# Mucosal-Associated Invariant T Cells Promote Atherosclerosis Through Monocyte-Driven Inflammation

**DOI:** 10.64898/2026.01.19.700259

**Authors:** Tobias Radecke, Thomas Nipoti, Louise Z. Wang, Nirmala Mouttoulingam, Marc Diedisheim, Mouna Chajadine, Emilie Bacquer, Coraline Heron, Paul Alayrac, Charles Caër, Ludivine Laurans, Camille Knosp, Pierre Liénart, Kenza Damache, Rida Al-Rifai, Stephane Camus, Mathilde Lemitre, Marie Piollet, Oliver Soehnlein, Pierre Julia, Jean-Marc Alsac, Salma El Batti, Franck Letourneur, Clément Cochain, Alain Tedgui, Hafid Ait-Oufella, Tristan Mirault, Ziad Mallat, Olivier Lantz, Jean-Sebastien Silvestre, Sophie Lotersztajn, Soraya Taleb

## Abstract

Mucosal-associated invariant T (MAIT) cells are unconventional T lymphocytes that may contribute to inflammatory responses, although their specific role in atherosclerosis remains poorly understood. In this study, we identified MAIT cells within human atherosclerotic plaques and found that they were significantly enriched among CD3⁺ T cells in plaques compared to matched peripheral blood samples. MAIT cells within plaques exhibited an activated phenotype and showed upregulation of genes associated with inflammation and cellular activation, compared to circulating MAIT cells from the same patients. Using murine models, we found that low-density lipoprotein receptor (Ldlr)^−/−^ mice carrying the CAST locus, which confers naturally higher frequencies of MAIT cells, displayed increased MAIT cell accumulation in both the liver and atherosclerotic plaques when fed a high-cholesterol diet. In contrast, MAIT cell-deficient Ldlr^−/−CAST^ MR1^−/−^ mice exhibited a reduced atherosclerotic burden, diminished liver fibrosis, smaller myocardial infarcts following coronary artery ligation, without significant changes in plasma cholesterol levels. These atheroprotective effects were accompanied by lower monocyte counts in the bone marrow and blood, as well as reduced plaque macrophage accumulation in the plaques. Furthermore, deletion of CCR2, which impairs monocyte mobilization, abrogated the pro-atherogenic effects of MAIT cells, indicating that MAIT-driven atherogenesis occurs through a monocyte-dependent mechanism. Taken together, these findings identify MAIT cells as active contributors to vascular inflammation and position them as potential therapeutic targets for atherosclerosis and its complications.

## Introduction

Cardiovascular diseases (CVD), including atherosclerosis, are among the leading causes of mortality worldwide, accounting for approximately 18 million deaths per year^1^. Atherosclerosis is closely linked to many comorbidities such as liver diseases^2^, and is a major complication that can lead to myocardial infarction (MI)^3^.

Atherosclerosis is an inflammatory disease of the vascular wall, triggered by the sub-endothelial accumulation of apoB-rich lipoproteins, including, low-density lipoprotein (LDL), and their subsequent oxidation. This process is controlled by both innate and adaptive immunity^4^. Innate immune cells, such as monocytes and macrophages, play a clear pro-atherogenic role^5^. Adaptive immunity also contributes significantly, with various T cell subsets exerting distinct effects. T helper (Th)1 cells, driven by the transcription factor T-box expressed in T cells (T-bet), promote atherosclerosis via interferon-gamma (IFN-γ). Th17 cells, regulated by Retinoic acid-related orphan receptor gamma t (RORγt), have conflicting results through interleukin-17 (IL-17), whereas T regulatory cells (Tregs) expressing the transcriptional factor forkhead box P3 (Foxp3) exert a protective effect^4,6,7^. CD8^+^ T cells also demonstrate a pro-atherogenic role, particularly through granzyme B production^8^.

Although our understanding of inflammation in atherosclerosis has advanced, the complex mechanisms by which innate and adaptive immune responses regulate its development and progression remain unclear. In particular, the role of mucosal-associated invariant T (MAIT) cells, a unique subset of unconventional T cells, is not yet fully defined in this context. These cells express a semi-invariant αβ T-cell receptor (Vα7.2-Jα33 in humans; Vα19-Jα33 in mice) and exhibit rapid, innate-like responses while retaining antigen-specific memory towards an ubiquitous bacterial antigen 5-(2-oxopropylideneamino)-6-d-ribitylaminouracil (5-OP-RU). This dual functionality positions them at the interface between innate and adaptive immunity, contributing not only to antimicrobial defence but also to inflammatory-mediated diseases^9^. Although abundant in human blood and mucosae, MAIT cells are relatively rare in murine organs^10^, challenging functional studies in mice. However, the development of CAST mice, which naturally exhibit a higher frequency of MAIT cells, due to a CAST genetic trait mapping to the TCR-α locus resulting in increased usage of distal Vα segments, including Vα19, provides a valuable model for characterizing their function^11^. MAIT cells recognize microbial-derived vitamin B2 (riboflavin) precursor presented by the non-classical MHC-related molecule (MR1) ^9^, which is crucial for MAIT development and function^12,13,14^. MAIT cell activation occurs via MR1-dependent TCR signaling or cytokine-mediated pathways, both of which contribute to their inflammatory and antimicrobial properties^12,15^.

Circulating MAIT cells have been reported to decrease in various chronic inflammatory diseases, including metabolic disorders^16,9^, and more recently, in patients with CVD^17,18^. However, the role of MAIT cells in CVD, particularly atherosclerosis, remains poorly understood. In this study, we investigated their involvement in atherosclerosis using B6-MAIT^CAST^ mice, which naturally have elevated MAIT cell numbers while maintaining normal development and function^19^. To assess the specific contribution of MAIT cells in atherosclerosis, we used both B6-MAIT^CAST^ and MAIT cell-deficient MR1^−/−^ B6-MAIT^CAST^ mice under proatherogenic background. Contrary to a recent study reporting a protective role for MAIT cells in promoting cholesterol removal^20^, our findings demonstrate that MAIT cells contribute to atherogenesis by modulating monocyte and macrophage number, independently of plasma cholesterol levels. We further investigated the presence of MAIT cells within human atherosclerotic plaques and conducted an in-depth characterization of their activation state and expression profile. Notably, MAIT cells within plaques displayed a higher activation state compared to MAIT cells in peripheral blood from the same patients.

## Results

### MAIT cells are enriched in human atherosclerotic carotid plaques and exhibit an activated phenotype

We detected MAIT cells in atherosclerotic plaques from patients undergoing carotid endarterectomy (clinical data in **Supp. Table 1**) using MR1 tetramers loaded with 5-OP-RU, whereas no staining was observed with the negative control MR1 tetramers^21^ loaded with 6-formylpterin (6-FP) (**Fig. 1A**). Quantification of MAIT cells in plaques revealed an increased proportion of these cells (expressed as a percentage of CD3⁺ T cells) compared to their counterparts in the peripheral blood of the same patients (**Fig. 1B, Supp. Fig 1A-B**). Moreover, their abundance in the plaque positively correlated with the proportion of MAIT cells in the peripheral blood of the same patients (**Supp. Fig 1C**). MAIT cells were in majority CD8^+^ in plaques (nearly 70%), but still significantly lower than what is observed in the peripheral blood (nearly 80%). The remaining cells were CD4^−^CD8^−^ double-negative MAIT cells, for which no significant differences were detected between the two compartments (**Fig 1C**). Moreover, MAIT cells within atherosclerotic plaques exhibited an activated phenotype, as evidenced by increased expression of the activation marker CD25 and the activation/residency marker CD69, as well as the exhaustion marker T cell immunoglobulin mucin-3 (TIM-3), alongside reduced expression of the anti-apoptotic marker (B-cell lymphoma 2) BCL-2, compared to circulating MAIT cells (**Fig. 1D**). These findings indicate that MAIT cells in plaques are more activated and consequently exhibit greater susceptibility to exhaustion and apoptosis. Regarding MAIT cell polarization into functional subsets (granzyme B–producing, IFN-γ–producing, and IL-17–producing MAIT cells), no significant differences were observed between circulating MAIT cells and those in plaques, except for a trend toward an increased frequency of IL-17–producing MAIT cells within plaques (**Supp. Fig. 1D-F**).

**Figure 1.**
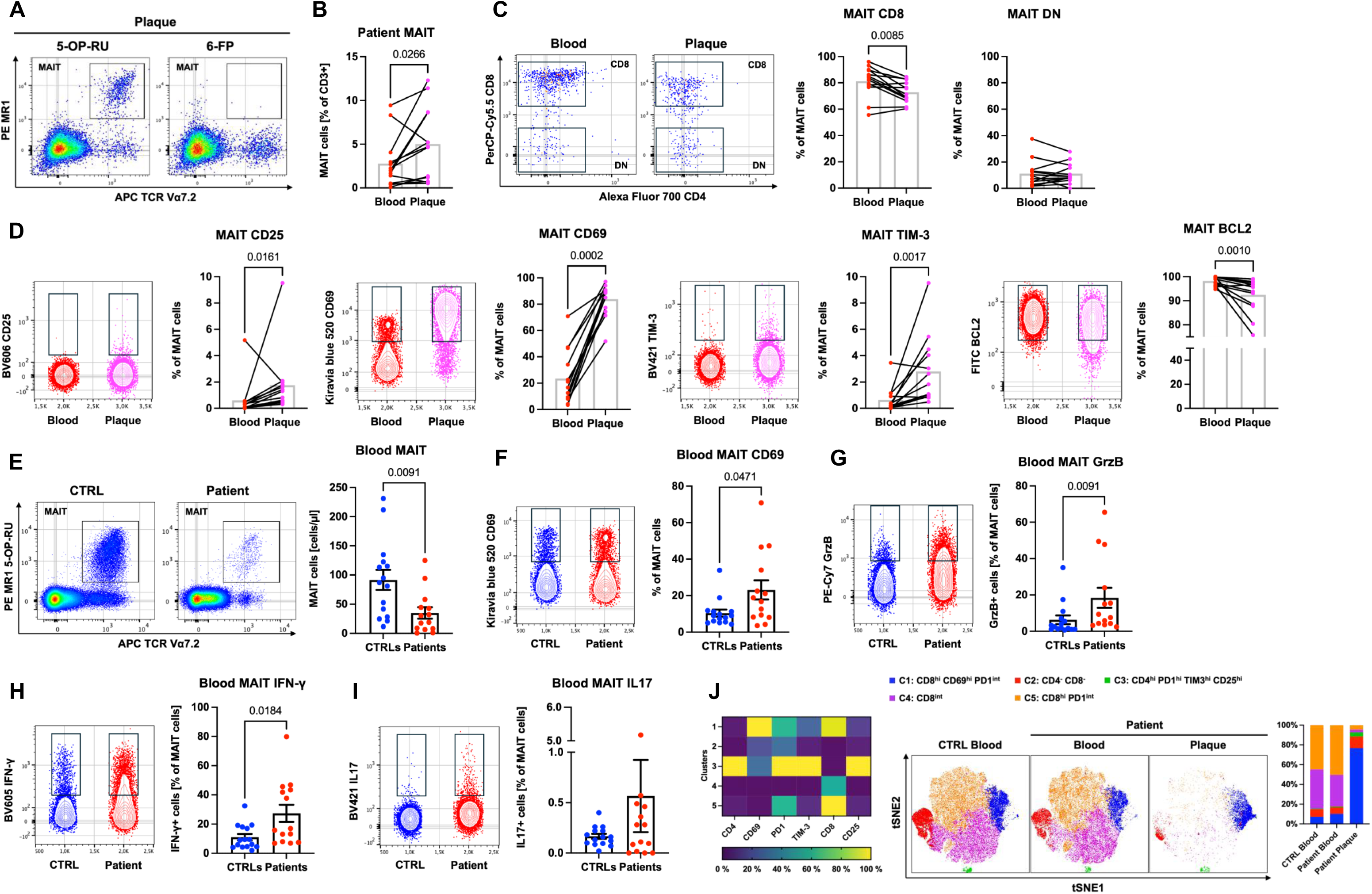
Presence of MAIT cells in human plaques exhibiting an activated state. **A.** Representative pseudocolor plots show MAIT cell staining in human atherosclerotic plaques from patients undergoing carotid endarterectomy using the 5-OP-RU tetramer and the negative control 6-FP tetramer. **B**. MAIT cell frequencies among CD3⁺ cells were quantified in PBMCs and plaques from the same patients (n = 13). **C**. Representative pseudocolor plots and quantification of CD8⁺ and double-negative (DN) CD4^−^CD8^−^ MAIT cell subsets in PBMCs and plaques from the same patients (n = 13). **D**. Representative contour plots and quantification of the percentages of CD25⁺, CD69⁺, TIM-3⁺, and BCL2⁺ MAIT cells in PBMCs and plaques from the same patients (n = 13). **E**. Representative pseudocolor plots and quantification of MAIT cell numbers in PBMCs from healthy controls (CTRLs, n = 15) and patients (n = 14). **F-H.** Representative contour plots and quantification of the percentages of CD69⁺, Granzyme B (GrzB⁺), and IFN-γ⁺ MAIT cells in PBMCs from healthy controls (CTRLs, n = 15) and patients (n = 14). **J.** An unbiased multi-dimensional analysis using X-Shift clustering identified five MAIT cell clusters; a heatmap shows the relative expression of six markers, and t-SNE plots illustrate the spatial distribution of these clusters. Data are presented as before–after plots or scatter plots with mean ± SEM. P-values were calculated using the Wilcoxon test (paired data) or Mann–Whitney test (unpaired data).

The increased proportion of MAIT cells within plaques could result from an enhanced recruitment of their counterparts in the peripheral blood. We therefore compared the number of MAIT cells in the peripheral blood between patients with CVD and healthy controls. In line with previous observations in chronic inflammatory diseases and recent reports in patients with CVD^17,18^, we found a significant reduction in circulating MAIT cells in patients with CVD compared to healthy donors (**Fig. 1E**). The majority of MAIT cells from peripheral blood mononuclear cells (PBMCs) were CD8^+^ (nearly 80%), with approximately 10% being double negative (CD8^−^CD4^−^) either in patients with CVD or in healthy controls (**Supp. Fig. 1G**). They were also more activated, as indicated by increased percentage of CD69 compared to controls (**Fig. 1F**). Geometric mean fluorescence intensity of the activation markers CD25 and CD69, and the exhaustion marker TIM-3 expressed in MAIT cells, negatively correlated with circulating MAIT cell numbers (**Supp. Fig. 1H**), suggesting that activation and/or exhaustion may also be involved in their reduced frequency in the blood. Moreover, MAIT cells (as % of CD3^+^ cells) in plaques were positively correlated with blood monocyte counts, and a similar trend was also observed between circulating MAIT cells and blood monocytes (**Supp. Fig. 1I**), suggesting a potential link between these cell populations.

In terms of effector molecule expression, circulating MAIT cells from patients with CVD exhibited higher levels of granzyme B (GrzB) and IFN-γ compared to controls, while IL-17 expression was not statistically different between the groups (**Fig. 1G-I**).

T-distributed stochastic neighbor embedding (t-SNE) analysis revealed an enrichment of CD8^high^ MAIT cell clusters expressing CD69 and intermediate levels of the exhaustion marker programmed cell death protein (PD)-1, in atherosclerotic plaques, compared to circulating MAIT cells from both patients and healthy controls (**Fig. 1J**). Taken together, these data suggest that MAIT cells within plaques are more activated and may be functionally distinct from their circulating counterparts.

To further characterize MAIT cells in plaques and blood, we sorted MAIT cells from PBMCs of healthy controls and from both PBMCs and plaques of the same patients with CVD, followed by bulk RNA sequencing. Principal component analysis (PCA) revealed distinct transcriptional profiles among the three different compartments (**Fig. 2A**). When comparing MAIT cells in PBMCs from healthy donors and patients, the top 50 differentially expressed genes were associated with immune cell activation and regulation, with up-regulation of SLAMF6, KLRG1, CCL4, and FKBP5 observed in patients, suggesting enhanced immune activation, cytotoxic potential, chemokine-mediated recruitment, and altered stress response pathways (**Fig. 2B**). Gene Ontology (GO) analysis revealed enrichment in pathways related to protein folding, immune activation, apoptosis, and stress responses (**Fig. 2C**). Moreover, when comparing MAIT cells from plaques to MAIT cells from the peripheral blood of the same patients, we observed a strong transcriptional shift with 625 genes upregulated and 463 downregulated. The top 50 differentially expressed genes were predominantly upregulated, except for the 50^th^ gene, and were strongly associated with inflammation (e.g. IFNγ, CCL4), stress responses (HSPA1A, HSP90AA1), regulation of apoptosis (MCL1, PMAIP1), and immune activation (CD69, TNFAIP3) (**Fig. 2D**). This highlights an activated, pro-inflammatory, and stress-adapted phenotype, consistent with findings from GO analysis (**Fig. 2E**). Taken together, these findings suggest that MAIT cells within atherosclerotic plaques undergo functional reprogramming to support local inflammation that may contribute to plaque progression.

**Figure 2.**
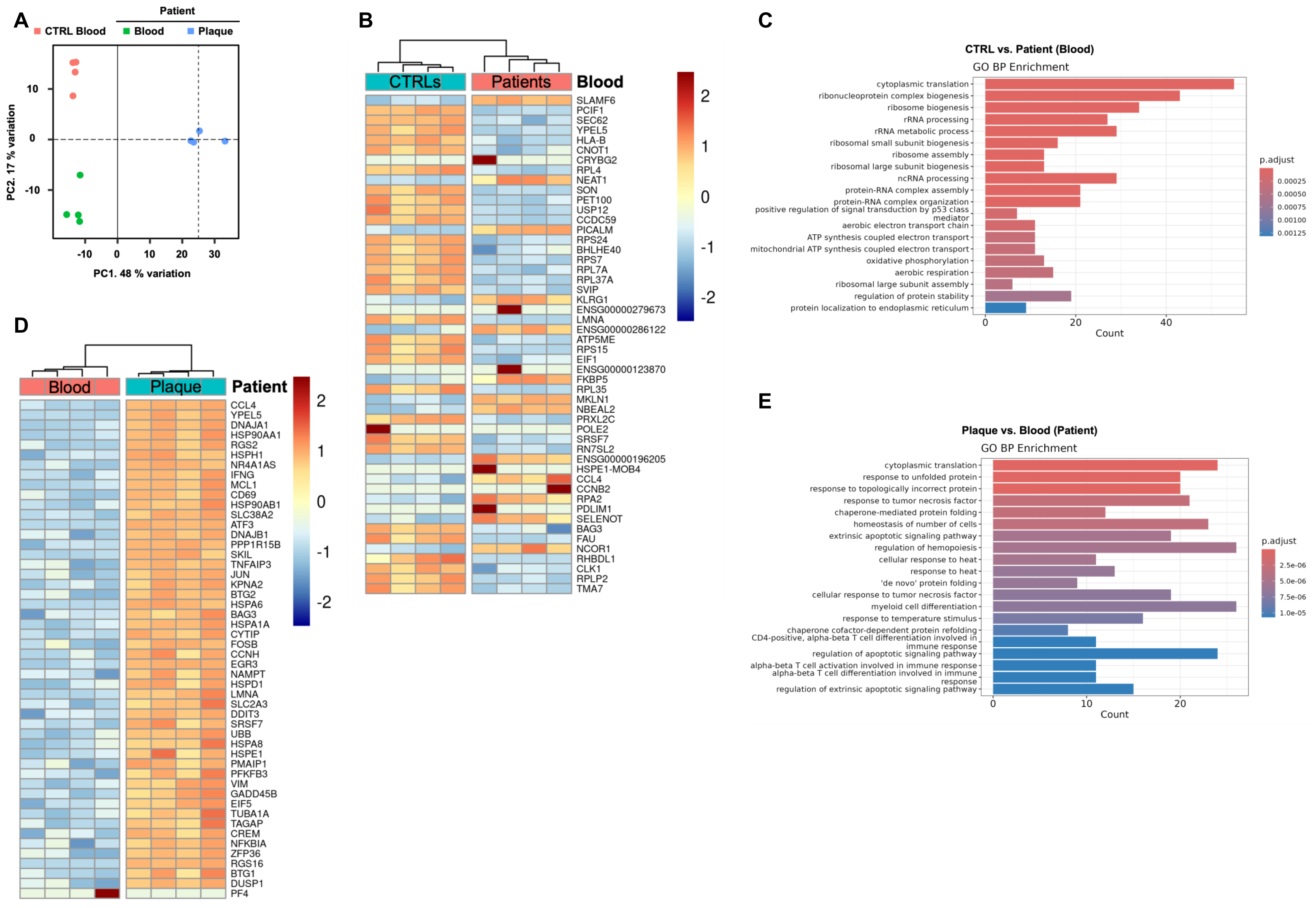
MAIT cells in human plaques exhibit an inflammatory profile. **A**. Principal component analysis (PCA) of MAIT cells isolated from healthy control (CTRL) blood, patient blood, and patient plaques (n = 4 per group) showing distinct clustering by sample origin. **B**. Heatmap of the top 50 differentially expressed genes between MAIT cells from CTRL blood and patient blood (n = 4 per group, z-score displayed). **C.** Gene Ontology (GO) enrichment analysis of genes differentially expressed between CTRL blood and patient blood (adjusted p-value < 0.001), revealing enrichment in immune and inflammatory pathways. **D.** Heatmap of the top 50 differentially expressed genes between MAIT cells from patient blood and patient plaque (n = 4 per group, z-score displayed). **E.** GO enrichment analysis of genes differentially expressed between patient blood and patient plaque (adjusted p-value < 0.001), highlighting pathways involved in inflammation and stress response.

### MAIT cells accumulate in plaques and exhibit activation markers in atherosclerotic mice

To investigate the role of MAIT cells in atherosclerosis, we used the low-density lipoprotein receptor-deficient (Ldlr^−/−^) mouse model, which is genetically prone to developing atherosclerosis when fed a high-cholesterol (HC) diet. Since MAIT cells are scarce in standard laboratory mice, we crossed B6-MAIT^CAST^ mice, which naturally exhibit higher MAIT cell frequencies^11^, with Ldlr^−/−^ mice (Ldlr^−/−CAST^). Using the MAIT staining detection strategy (**Supp. Fig. 2A**), we found that Ldlr^−/−CAST^ mice exhibited a fivefold increase in MAIT cell numbers than Ldlr^−/−^ littermates (**Fig. 3A**) in the liver, a well-known MAIT cell–enriched organ^9^. To study dynamic changes in MAIT cells during atherosclerosis, Ldlr^⁻/⁻CAST^ mice were fed either a chow diet (CD) or HC diet for a short duration (1 week) or an extended period (8 weeks) (**Fig. 3B**). Following 1 week of HC diet, circulating MAIT cell numbers, which were initially low, showed a decreasing trend (p = 0.0556; **Fig. 3C**), while no significant changes were observed after 8 weeks. (**Supp. Fig. 2B**). In the spleen, MAIT cell numbers decreased after 8 weeks of HC diet (**Fig. 3D**). In contrast, MAIT cells increased in the liver after both 1 week (**Supp. Fig. 2C**) and 8 weeks of HC feeding (**Fig. 3E**). We then aimed to assess whether MAIT cells migrated to atherosclerotic plaques, particularly in the aorta. Using MR1 tetramer loaded with 5-OP-RU, we detected a small number of MAIT cells in the aorta, with their numbers increasing under HC diet (**Fig. 3F**). After 8 weeks of HC diet, most MAIT cells in the liver and spleen exhibited a double-negative (CD4^−^CD8^−^) phenotype, whereas MAIT cells in the aorta were almost entirely CD4^−^CD8^−^. The overall proportions of CD4⁺, CD8⁺, and double-negative MAIT cell subsets remained largely unchanged between the HC and CD groups, with the exception of a modest increase in the frequency of CD8⁺ MAIT cells in the spleen of HC diet-fed mice (**Supp. Fig. 2D–E**; **Fig. 3G**). Taken together, these results suggest that under the HC diet, MAIT cells decrease in the blood and spleen, potentially due to their migration to peripheral tissues such as the liver and aorta.

**Figure 3.**
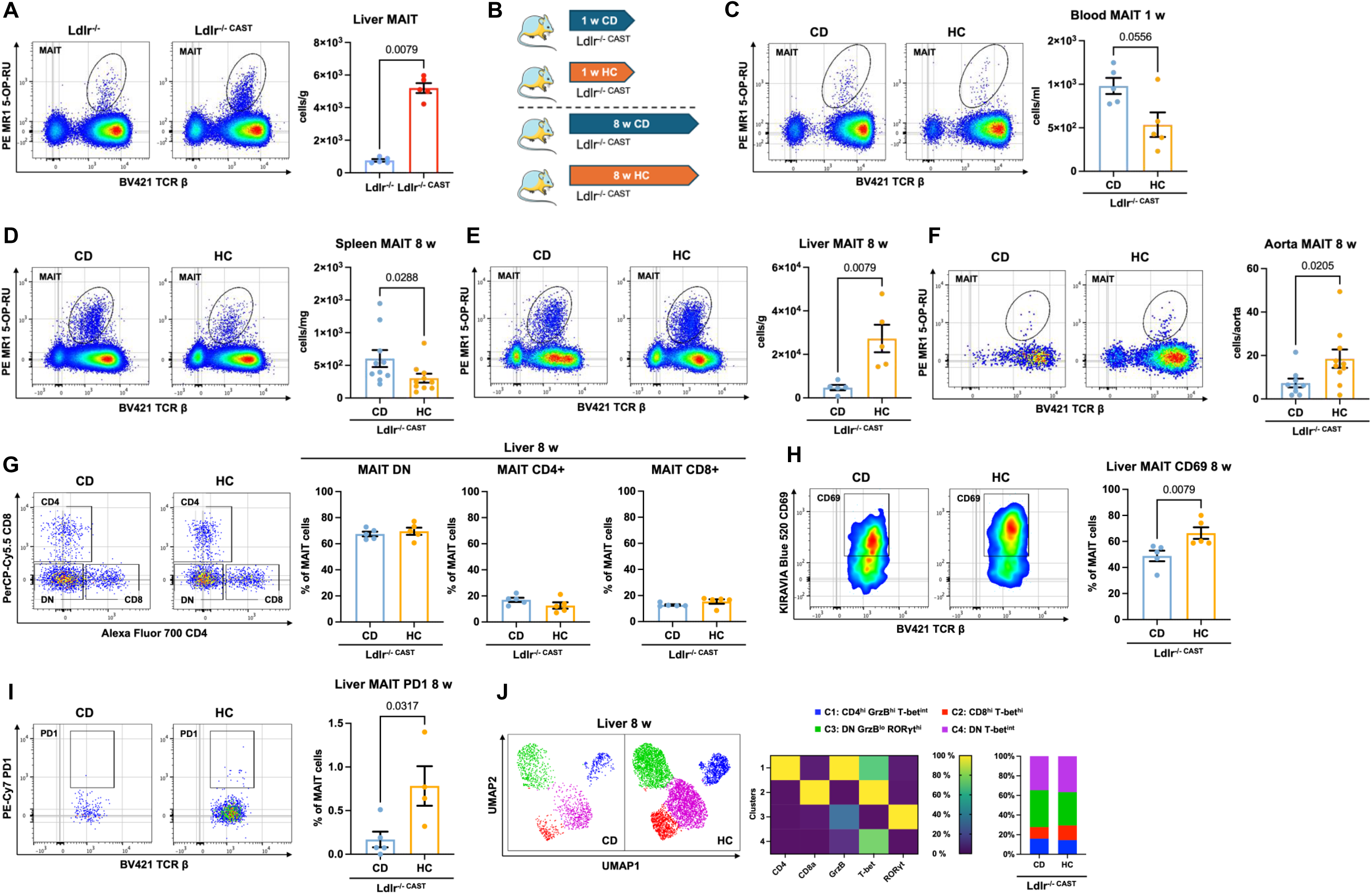
MAIT cells exhibit an activated profile under an atherogenic diet in mice. **A**. Representative pseudocolor plots and quantification of MAIT cell numbers per gram of tissue, based on 5-OP-RU tetramer staining of liver extracts from female Ldlr^−/−^ and Ldlr^−/−CAST^ mice fed a chow diet (CD) (n = 5 per group). **B**. Schematic representation of the experimental design. **C.** Representative pseudocolor plots and quantification of MAIT cell numbers in the blood of male Ldlr^−/−CAST^ mice fed a CD or HC diet for 1 week (n = 5 per group). **D.** Representative pseudocolor plots and quantification of MAIT cell numbers per gram of spleen tissue from male Ldlr^−/−CAST^ mice fed a CD or HC diet for 8 weeks (n = 10 per group, a pool of two independent experiments). **E**. Representative pseudocolor plots and quantification of MAIT cell numbers per gram of liver tissue from male Ldlr^−/−CAST^ mice fed a CD or HC diet for 8 weeks (n = 5 per group, replicated in 2 independent experiments). **F.** Representative pseudocolor plots and quantification of MAIT cell numbers per completely digested aorta from male Ldlr^−/−CAST^ mice fed a CD or HC diet for 8 weeks (n = 9-10 per group, a pool of two independent experiments). **G.** Representative pseudocolor plots and quantification of CD4⁺, CD8⁺, and double-negative (DN) CD4^−^CD8^−^ MAIT cells from the livers of male Ldlr^−/−CAST^ mice fed a CD or HC diet for 8 weeks (n = 5 per group, replicated in 2 independent experiments). **H-I**. Representative pseudocolor plots and quantification of the percentages of CD69⁺ and PD-1⁺ MAIT cells from livers of male Ldlr^−/−CAST^ mice fed a CD or HC diet for 8 weeks (n = 4-5 per group, replicated in 2 independent experiments). **J.** Unbiased multi-dimensional analysis of MAIT cells from livers of male Ldlr^−/−CAST^ mice fed a CD or HC diet for 8 weeks (n = 5 per group), using X-Shift clustering and UMAP visualization. Heatmap depicts relative expression levels of five markers across the four identified clusters. Individual data are presented as scatter plots with mean and SEM. P-values were calculated using the Mann-Whitney test.

MAIT cells in the liver after 1 week of HC feeding showed an increase in CD69 expression (**Supp. Fig. 3A-B**), which was also observed after 8 weeks of HC feeding in both the spleen and liver (**Supp. Fig. 3C**, **Fig. 3H**). This rise in MAIT cell activation was associated with a slight increase in the exhaustion marker PD-1 (%) in both the liver and spleen after 8 weeks of HC feeding (**Fig. 3I**, **Supp. Fig. 3D**). In aortic MAIT cells, no significant changes in CD69 expression were observed after 8 weeks of HC diet compared to CD (**Supp. Fig. 3E**). Polarization of MAIT cells toward MAIT1 (T-bet⁺) and MAIT17 (RORγt⁺) subsets, with MAIT17 representing over 90% of aortic MAIT cells, showed no significant differences between dietary conditions, based on FACS analysis of T-bet⁺ and RORγt⁺ expression as a percentage of total MAIT cells (**Supp. Fig. 3F**). Similarly, in the spleen, MAIT17 (RORγt⁺) cells accounted for nearly 40% of the total MAIT population, and the proportion of both MAIT17 (RORγt⁺) or MAIT1 (T-bet⁺) cells remained unchanged between CD and HC diet groups (**Supp. Fig. 3G**). In the liver, a significant decrease in the percentage of MAIT1 (T-bet⁺) cells was observed under HC diet, whereas MAIT17 (RORγt⁺) cells were not affected (**Supp. Fig. 3H**). Uniform Manifold Approximation and Projection (UMAP) analysis in the liver revealed that, in addition to the predominant CD4^−^CD8^−^ RORγt^high^ subset, a distinct population of CD4^−^CD8^−^ MAIT cells expressing intermediate levels of T-bet was also present (**Fig. 3J**). Taken together, these data show a marked accumulation of MAIT17 cells in inflammatory tissues such as the liver and atherosclerotic aortas, accompanied by enhanced activation of these cells in both the spleen and liver.

### MAIT cells exacerbate atherosclerosis and its associated complications

To investigate the role of MAIT cells in atherosclerosis, we first performed bone marrow transplantation (BMT) in Ldlr^−/−^ mice. The mice were irradiated with 9.5 Gray and transplanted with bone marrow from either C57BL/6 or B6-MAIT^CAST^ mice (**Fig. 4A**). As shown in **Fig. 4B–4C** and **Supp. Fig. 4A**, MAIT supplementation in mice receiving B6-MAIT^CAST^ bone marrow showed an increase in plaque size in both the aortic root and thoracic aorta. Although a slight increase in plasma cholesterol levels was observed in mice transplanted with B6-MAIT^CAST^ bone marrow, this is unlikely to have had a significant impact on atherosclerosis, as no significant correlation was found between plasma cholesterol levels and plaque size in either the aortic root or the thoracic aorta (**Supp. Fig. 4B–4C**). We also fed Ldlr^−/−CAST^ mice (exhibiting high levels of MAIT cells), their littermate Ldlr^−/−^ mice (with low MAIT cells), and Ldlr^−/−CAST^ MR1^−/−^ mice (MAIT cell-deficient) a HC diet for 8 weeks, in both female and male mice (**Fig. 4D**). As shown in **Fig. 4E–G**, a high MAIT cell number in Ldlr^−/−CAST^ female mice was associated with increased plaque size (% of total thoracic aorta) compared to Ldlr^−/−CAST^ MR1^−/−^ mice or Ldlr^−/−^ mice, without significant changes in plasma cholesterol levels. A similar trend was observed in males, with Ldlr^−/−CAST^ mice showing a significant increase in plaque size compared with Ldlr^−/−^ mice (p-value = 0.0016), while no significant difference was detected between Ldlr^−/−^ and Ldlr^−/−CAST^ MR1^−/−^ mice (p-value = 0.3326), and plasma cholesterol levels remained comparable across groups (**Supp. Fig. 4D–F**).

**Figure 4:**
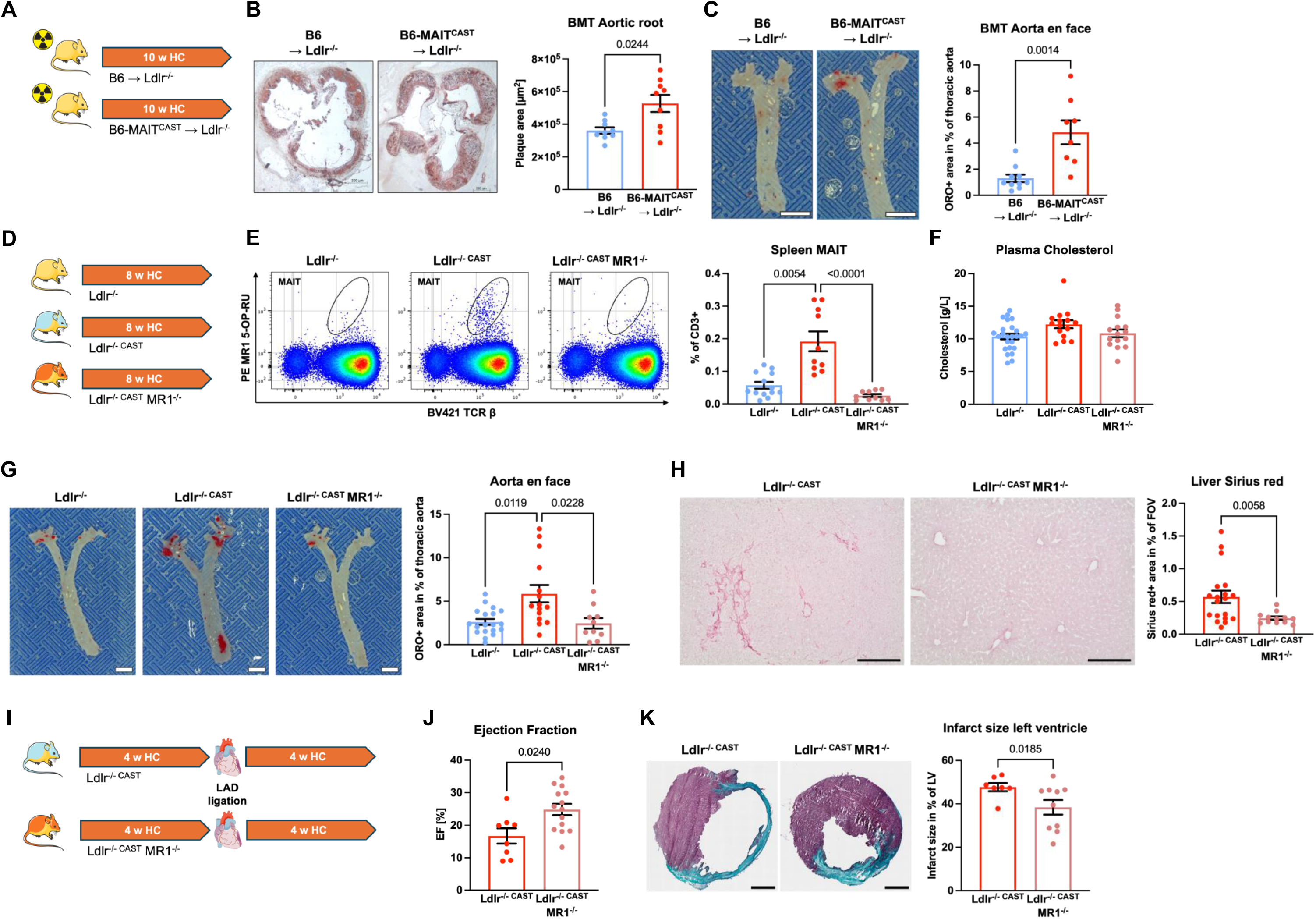
Absence of MAIT cells relieved atherosclerosis and its associated complications. **A.** Schematic representation of the experimental design. **B.** Representative images and quantification of Oil Red O-stained aortic root sections from irradiated female Ldlr^−/−^ mice undergoing bone marrow transplantation (BMT) that received bone marrow from female B6 or B6-MAIT^CAST^ donors, followed by a 10-week high-cholesterol (HC) diet (n = 9 per group, scale bar, 200 µm). **C.** Representative images and quantification of Oil Red O-stained thoracic aortas from irradiated female Ldlr^−/−^ mice receiving bone marrow from female B6 or B6-MAIT^CAST^ donors after a 10-week HC diet (scale bar, 2 mm). **D.** Schematic representation of the experimental design. **E.** Representative pseudocolor plots and quantification of MAIT cells stained with 5-OP-RU tetramer from spleens of female Ldlr^−/−^ (n = 13), Ldlr^−/−CAST^ (n = 10), and Ldlr^−/−CAST^ MR1^−/−^ (n = 10) mice fed HC diet for 8 weeks (pool of three independent experiments). **F.** Plasma cholesterol levels in female Ldlr^−/−^ (n = 25), Ldlr^−/−CAST^ (n = 15), and Ldlr^−/−CAST^ MR1^−/−^ (n = 15) mice after 8 weeks on an HC diet (pool of three independent experiments). **G.** Representative images and quantification of Oil Red O-stained thoracic aortas from female Ldlr^−/−^ (n = 19), Ldlr^−/−CAST^ (n = 15), and Ldlr^−/−CAST^ MR1^−/−^ (n = 10) mice on an HC diet for 8 weeks (scale bar, 2 mm, pool of three independent experiments). **H.** Representative images and quantification of Picro-Sirius Red-stained liver sections from female Ldlr^−/−CAST^ (n = 19) and Ldlr^−/−CAST^ MR1^−/−^ (n = 12) mice on an HC diet for 8 weeks (scale bar, 200 µm, pool of two independent experiments). **I.** Schematic representation of the experimental design. **J.** Quantification of ejection fraction in male Ldlr^−/−CAST^ (n = 8) and Ldlr^−/−CAST^ MR1^−/−^ (n = 14) mice fed a HC diet for 8 weeks and undergoing left anterior descending (LAD) coronary artery ligation. **K.** Representative images and quantification of Masson Trichrome-stained heart sections from male Ldlr^−/−CAST^ and Ldlr^−/−CAST^ MR1^−/−^ mice fed a HC diet and LAD ligation (scale bar, 200 µm). Individual data are presented as scatter plots with mean ± SEM. Mann-Whitney tests were used for two-group comparisons; Kruskal-Wallis followed by Dunn’s multiple comparisons test was used for more than two groups.

We also examined atherosclerotic plaque development in the aortic root and found that female Ldlr^−/−CAST^ MR1^−/−^ mice exhibited reduced plaque size compared to Ldlr^−/−^ mice, whereas no significant differences were observed in male mice (**Supp. Fig. 5A–B**). The reduction in plaque size in the aortic root observed in female Ldlr^−/−CAST^ MR1^−/−^ mice was associated with a significant decrease in macrophage accumulation, as assessed by MOMA-2 staining, with no significant changes in T cell (CD3⁺) numbers, as well as a reduction in fibrosis, assessed by Sirius Red staining (**Supp. Fig. 5C–5E**). The more pronounced female phenotype may be explained, at least in part, by increased MAIT-cell activation compared with males, as supported by the higher CD25 expression observed in female mice **(Supp. Fig. 5F**). Following a prolonged period of HC feeding (14 weeks), no differences in plaque size were observed in either the thoracic aorta or aortic root between male or female Ldlr^−/−CAST^ mice and Ldlr^−/−CAST^ MR1^−/−^ mice (**Supp Fig. 6A-E**), suggesting that MAIT cells exert a pro-atherogenic effect primarily during the early stages of atherosclerosis.

**Figure 5.**
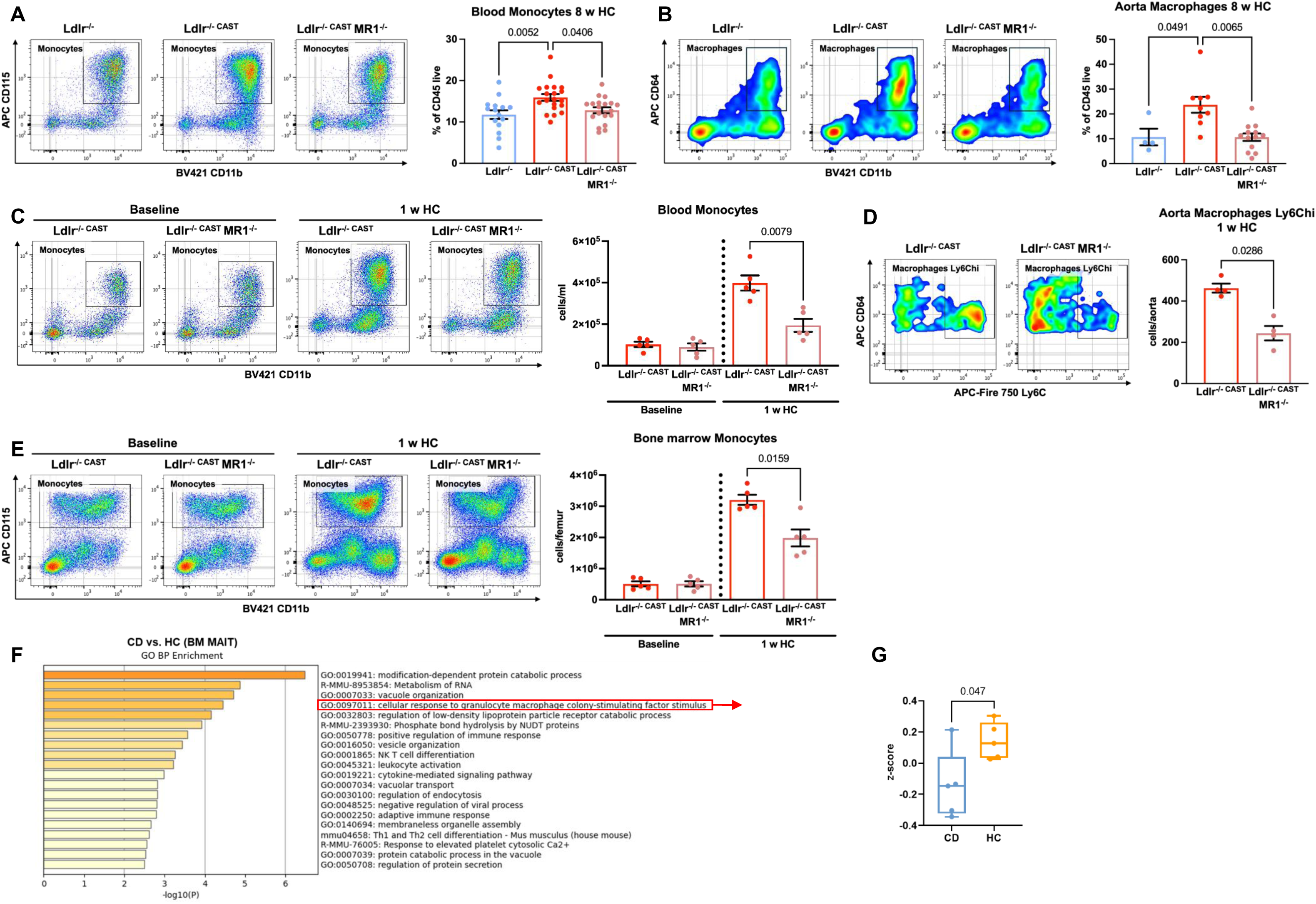
Absence of MAIT cells is associated with reduced bone marrow/blood monocyte and aortic macrophage numbers. **A.** Representative pseudocolor plots and quantification of blood monocytes from female Ldlr^−/−^ (n = 15), Ldlr^−/−CAST^ (n = 20), and Ldlr^−/−CAST^ MR1^−/−^ (n = 19) mice fed a high-cholesterol (HC) diet for 8 weeks (pool of four independent experiments). **B.** Representative pseudocolor plots and quantification of aortic macrophages from female Ldlr^−/−^ (n = 4), Ldlr^−/−CAST^ (n = 9), and Ldlr^−/−CAST^ MR1^−/−^ (n = 13) mice after 8 weeks on a HC diet (pool of two independent experiments). **C.** Representative pseudocolor plots and quantification of blood monocytes (CD11b^+^ CD115^+^) at baseline and after 1 week of HC (n = 5 per group). **D.** Aortic Ly6C^hi^ macrophages (CD11b^+^ CD64^+^) from female Ldlr^−/−CAST^ and Ldlr^−/−CAST^ MR1^−/−^mice (n = 4 per group) after 1 week of HC diet. **E.** Representative pseudocolor plots and quantification of bone marrow monocytes (CD11b^+^ CD115^+^) at baseline and after 1 week of HC (n = 5 per group). **F.-G.** Bulk RNA sequencing of MAIT cells sorted from bone marrow in Ldlr^−/−CAST^ mice fed a CD or a HC diet for 1 week (n = 5 per group). Gene Ontology (GO) enrichment analysis of genes differentially expressed (p-value < 0.02) revealing enrichment in cellular response to granulocyte macrophage colony-stimulating factor stimulus pathway (highlighted in red) (F) and corresponding z-score of this pathway between the two diet conditions (G). The Individual data are presented as scatter plots, with mean ± SEM. The Mann-Whitney test was used for comparisons between two groups, and the Kruskal-Wallis test followed by Dunn’s multiple comparisons was used for three or more groups. Dashed lines indicate that the experiments were not done at the same time and the Mann-Whitney test was used for comparison between the groups.

**Figure 6.**
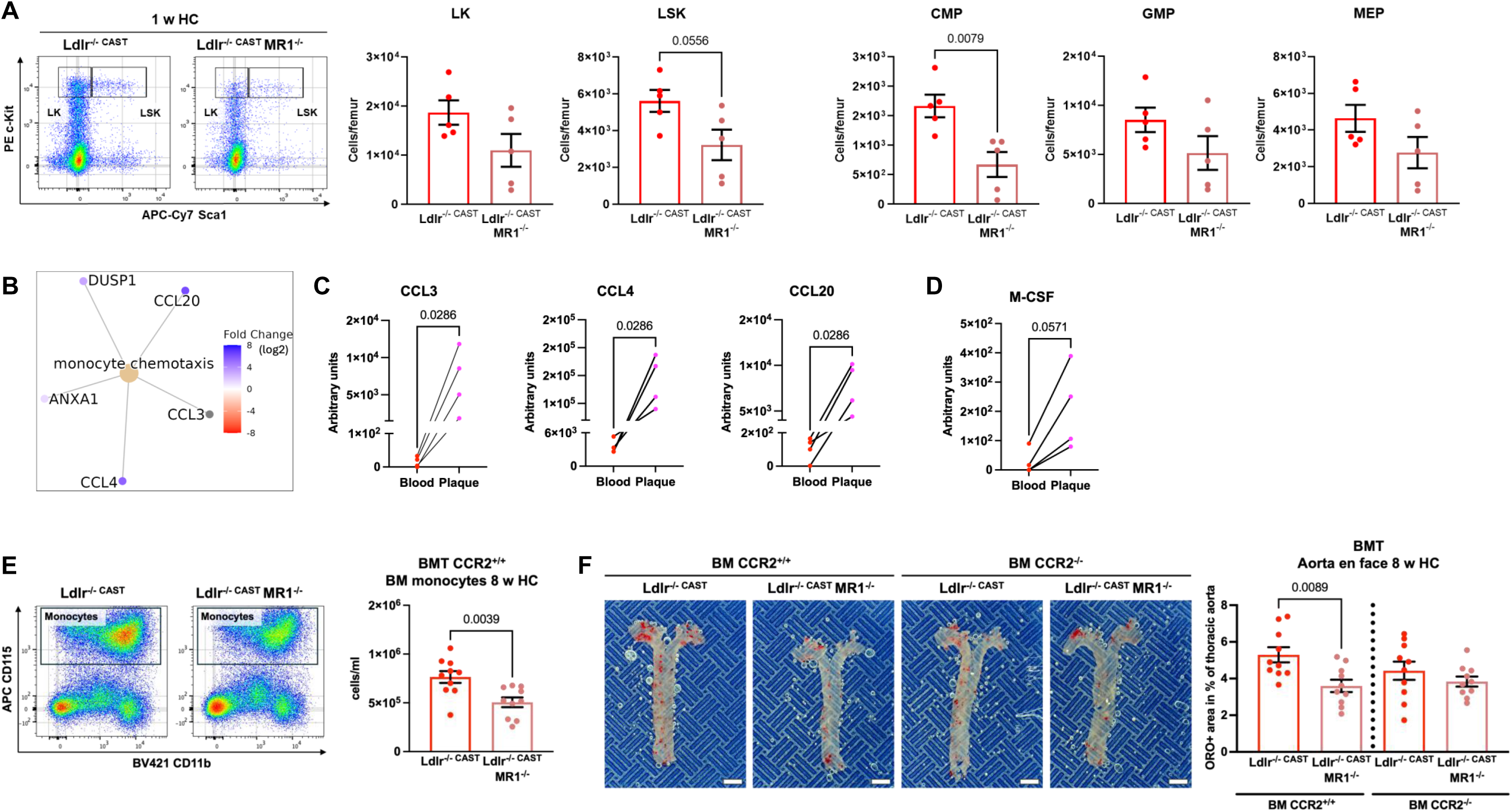
Absence of MAIT cells alleviates atherosclerosis by reducing monocyte output through altered bone marrow myelopoiesis. **A.** Representative pseudocolor plots and quantification of bone marrow LK (Lin^−^ c-Kit^high^ Sca1^low^), LSK (Lin^−^c-Kit^high^ Sca1^high^), CMP (Lin^−^c-Kit^high^ Sca1^low^ CD34^high^ CD16/32^low^), GMP (Lin^−^ c-Kit^high^ Sca1^low^ CD34^high^ CD16/32^high^) and MEP (Lin^−^ c-Kit^high^ Sca1^low^ CD34^low^ CD16/32^low^) progenitors from female Ldlr^−/−CAST^ and Ldlr^−/−CAST^ MR1^−/−^ mice fed a high-cholesterol (HC) diet for 1 week. The lineage (Lin) cocktail is defined as a mixture of anti-CD3ε PE-Cy7, anti-Ter-119 PE-Cy7, anti-CD45R PE-Cy7, and anti-Gr1 PE-Cy7 antibodies (n = 5 per group, replicated in 2 independent experiments). **B.-D.** Bulk RNA sequencing of MAIT cells sorted from human PBMCs and atherosclerotic plaques from the same patients (n = 4). Gene ontology (GO) analysis of differentially expressed genes revealed enrichment in monocyte chemotaxis–related pathways (B). Quantification of monocyte-attracting chemokines (CCL3, CCL4, CCL20) (C) and M-CSF (D) expression in plaque- and PBMC-derived MAIT cells is shown. **E.** Representative pseudocolor plots and quantification of bone marrow monocytes from irradiated female Ldlr^−/−CAST^ and Ldlr^−/−CAST^ MR1^−/−^ mice that undergo bone marrow transplantation (BMT), received bone marrow from female CCR2^+/+^ donors, followed by an 8-week HC diet (n = 10 per group). **F.** Representative images and quantification of Oil Red O-stained thoracic aortas from irradiated female Ldlr^−/−CAST^ and Ldlr^−/−CAST^ MR1^−/−^ mice that received bone marrow from CCR2^+/+^ or CCR2^−/−^ female donors and were fed a HC diet for 8 weeks (n = 10 per group; scale bar, 2 mm). Individual data are presented as before-after or scatter plots, with mean ± SEM. The Mann-Whitney test was used for comparisons between two groups. Dashed lines indicate that the experiments were not done at the same time and the Mann-Whitney test was used for comparison.

Liver fibrosis is associated with poorer liver disease outcomes and atheroslerosis^2^ and MAIT cells have been shown to exert deleterious, pro-fibrotic role^16,22^. Interestingly, Ldlr^−/−CAST^ MR1^−/−^ mice exhibited reduced liver fibrosis compared to Ldlr^−/−CAST^ mice (**Fig. 4H**). MI is recognized as one of the main clinical manifestations of atherosclerosis^3^. To investigate whether MAIT cell-deficient mice, which are protected against atherosclerosis, exhibit reduced susceptibility to MI, we induced MI by permanent ligation of the left coronary artery in Ldlr^−/−CAST^ and Ldlr^−/−CAST^ MR1^−/−^ mice fed a HC diet for 4 weeks, followed by a 4-week post-MI period (**Fig. 4I**). Ldlr^−/−CAST^ MR1^−/−^ mice, compared to Ldlr^−/−CAST^ mice subjected to both MI and atherosclerosis, continued to exhibit reduced atherosclerosis burden (**Supp. Fig. 6F–G**). In addition, a significant positive correlation was observed between plaque size and liver fibrosis in these mice (**Supp. Fig. 6H**). Interestingly, Ldlr^−/−CAST^ MR1^−/−^ compared to Ldlr^−/−CAST^ mice showed improved cardiac function, as evidenced by a higher ejection fraction and smaller infarct size assessed by Masson’s Trichrome staining (**Fig. 4J-K**). Taken together, these results demonstrate that MAIT cell deficiency confers protection not only against atherosclerosis but also against its associated complications, liver fibrosis and MI.

### MAIT cells exert a pro-atherogenic role through monocytes/macrophages

We then sought to elucidate the mechanisms underlying the pro-atherogenic role of MAIT cells. Given their function as a bridge between innate and adaptive immunity, we focused our analysis on changes in immune cell subsets within the blood, spleen, and aorta.

After 8 weeks of HC diet feeding, circulating monocyte levels (CD115^+^) were elevated in Ldlr^−/−CAST^ mice compared to both Ldlr^−/−^ and Ldlr^−/−CAST^ MR1^−/−^ mice (**Fig. 5A**). This increase encompassed both classical monocytes (Ly6C^high^) and non-classical monocytes (Ly6C^low^) (**Supp. Fig. 7** and **8A**). In contrast, blood levels of other immune cells, including neutrophils (Ly6G^+^), CD4^+^ and CD8^+^ T cells, and B cells (CD19^+^), did not differ significantly between the groups (**Supp. Fig. 7 and 8B**). In the aorta, macrophage (CD64^+^) numbers were increased in Ldlr^−/−CAST^ compared to Ldlr^−/−^ or Ldlr^−/−CAST^ MR1^−/−^ mice (**Fig. 5B**), whereas neutrophils (Ly6G^+^), CD4^+^ and CD8^+^ T cells, and B cells (CD19^+^) showed no significant differences (**Supp. Fig. 8C**). Similarly, in the spleen, total CD4⁺ T-cell numbers and their polarization into Th1 (T-bet⁺), Th17 (RORγt⁺), and Treg (Foxp3⁺) subsets (expressed as a percentage of CD4⁺ cells), as well as the frequencies of monocytes, neutrophils and macrophages, were comparable across groups (**Supp. Fig. 8D–E**).

Monocytosis, which occurs within a few days of HC diet feeding and has been shown to play a pro-atherogenic role^23^, prompted us to assess monocyte levels at baseline and after 1 week of HC feeding period. As shown in **Fig. 5C**, a fourfold increase in blood monocytes was observed in Ldlr^−/−CAST^ mice, while the increase was more modest, approximately twofold, in Ldlr^−/−CAST^ MR1^−/−^ mice. After 1 week of HC diet, a reduction in aortic macrophages derived from classical monocytes (CD64^+^ Ly6C^high^) was also noted in Ldlr^−/−CAST^ MR1^−/−^ mice compared to Ldlr^−/−CAST^ mice (**Fig. 5D**).

The lower levels of both circulating monocytes and aortic macrophages in mice lacking MAIT cells were also associated with a less pronounced increase in the bone marrow monocyte numbers after 1 week of HC diet compared to the baseline (**Fig. 5E**), suggesting that MAIT cells may influence monocyte dynamics within bone marrow. Supporting this hypothesis, MAIT cells were detected in the bone marrow using MR1 tetramers loaded with 5-OP-RU and displayed activation markers, including increased frequency of CD25⁺ and CD69⁺ cells, particularly after 8 weeks of HC diet (**Supp. Fig. 9A-E**).

To further investigate how MAIT cells may influence monocytes in the bone marrow during the early stages of atherosclerosis, we isolated MAIT cells using MR1 tetramers and performed bulk RNA sequencing on bone-marrow-sorted MAIT cells from Ldlr^−/−CAST^ mice fed either a CD or a HC for 1 week. Under HC conditions, among the top significantly enriched biological processes compared with CD (p-value < 0.02), MAIT cells exhibited pathways associated with immune activation, including leukocyte activation, immune effector response, and positive regulation of the immune response (**Fig. 5F**). Notably, the myelopoiesis-related pathway was enriched, such as the cellular response to granulocyte–macrophage colony-stimulating factor stimulus (**Fig. 5F**). This pathway was significantly elevated in bone-marrow MAIT cells from HC-fed mice compared with CD, as indicated by higher mean z-scores and increased ssGSEA enrichment scores (**Fig. 5G** and **Supp. Fig. 9F**). Within this pathway, several genes, including Irak2, Tlr1, Il18rap, Cd38, Txk, and Prkcb, were upregulated (**Supp. Fig. 9G**), reflecting an activated inflammatory state that may promote myelopoiesis and bias myeloid cell development.

To further investigate whether the smaller increase in bone-marrow monocytes observed in the absence of MAIT cells reflected impaired myelopoiesis, we analyzed upstream myeloid progenitor populations. After 1 week of HC feeding, Ldlr^−/−CAST^ MR1^−/−^ mice displayed significantly fewer common myeloid progenitors (CMP; Lin^−^ c-Kit^high^ Sca1^low^ CD34^high^ CD16/32^low^), reduced granulocyte–macrophage progenitors (GMP; Lin^−^ c-Kit^high^ Sca1^low^ CD16/32^high^ CD34^high^), decreased frequencies of LSK cells (Lin^−^c-Kit^high^ Sca1^high^), and a reduction in hematopoietic stem cells (HSC; Lin^−^ c-Kit^high^ Sca1^high^ CD135^low^ CD48^low^), particularly within the short-term HSC subset (ST-HSC; Lin^−^ c-Kit^high^ Sca1^high^ CD135^low^ CD48^low^ CD150^−^). In contrast, no significant differences between genotypes were observed under CD conditions (**Fig. 6A; Supp. Fig. 10–11**). Together, these data support a MAIT cell–dependent regulation of early myeloid development, highlighting a potential cross-talk between MAIT cells and the monocyte/macrophage lineage within the bone marrow.

MAIT cells are known to produce a range of chemokines (e.g., CCL3, CCL4) and growth factors (e.g. Granulocyte-Macrophage Colony-Stimulating Factor; GM-CSF^24^), which can strongly influence monocyte function and recruitment^25,26^. Consistently, we found that human MAIT cells isolated from atherosclerotic plaques of patients with CVD, which were more activated, showed upregulation of genes involved in monocyte chemotaxis (p-value = 0.0061), including chemokines such as CCL3, CCL4, and CCL20, compared to matched circulating MAIT cells from the same patients (**Fig. 6B-C**). Macrophage colony-stimulating factor (M-CSF) was also upregulated in MAIT cells from atherosclerotic plaques compared to circulating MAIT cells of the same patients (**Fig. 6D**). Taken together, these data suggest that MAIT cells become activated within inflammatory environments and contribute to atherosclerosis by producing factors that promote monocyte/macrophage recruitment and activation.

We then sought to determine whether the pro-atherogenic effects of MAIT cells depend on monocytes and macrophages *in vivo*. To investigate this, we performed BMT in Ldlr^−/−CAST^ and Ldlr^−/−CAST^ MR1^−/−^mice using either CCR2-deficient bone marrow (CCR2^−/−^), which impairs monocyte egress from the bone marrow into the circulation^27^, or CCR2-sufficient bone marrow (CCR2^+/+^, **Supp. Fig. 12A**). As shown in **Fig. 6E**, monocyte numbers in the bone marrow were significantly reduced in Ldlr^−/−CAST^ MR1^−/−^ mice receiving CCR2^+/+^ bone marrow compared to Ldlr^−/−CAST^ mice transplanted with the same bone marrow.

Notably, MAIT cell deficiency reduced atherosclerosis in CCR2^+/+^ bone-marrow–reconstituted mice, whereas plaque size was similarly reduced in CCR2^−/−^ chimeras, which showed low circulating monocyte levels irrespective of MAIT cell status (**Fig. 6F** and **Supp. Fig. 12B-C**). The lack of differences in circulating monocyte levels between MAIT-deficient and control mice under CCR2^+/+^ bone-marrow reconstitution may reflect dynamic monocyte trafficking from the blood into peripheral tissues. These effects occurred without significant differences in plasma cholesterol levels between groups and in the absence of any correlation between plaque size and circulating cholesterol (**Supp. Fig. 12D–E**). Moreover, in mice reconstituted with CCR2^+/+^ bone marrow, the reduction in bone-marrow monocytes observed in the absence of MAIT cells was accompanied by a trend toward decreased macrophages (MOMA-2⁺), particularly within Oil Red O–positive plaque areas (**Supp. Fig. 12F**). These results suggest that MAIT cells aggravate atherosclerosis through effects on monocytes and macrophages.

In conclusion, our findings demonstrate that MAIT cells become activated under atherogenic high-cholesterol conditions, undergoing significant phenotyping changes. This activation promotes an increase in monocyte and macrophage numbers, particularly within atherosclerotic plaques, thereby driving a pro-atherogenic effect independent of plasma cholesterol levels.

## Discussion

While MAIT cells have been previously studied in the context of chronic inflammation, including metabolic and liver diseases^16,9^ and more recently in CVD^17,18,20^, their presence and functional characteristics within human atherosclerotic plaques remain largely unexplored. In this study, we identified MAIT cells using MR1 tetramers loaded with 5-OP-RU within human plaques and found that they are more enriched in plaques than in the blood of the same patients with CVD. Furthermore, there was a negative correlation between MAIT cell number in peripheral blood and their proportion within atherosclerotic plaques from the same patients. Although the significance of this observation is not yet fully understood, it may suggest that MAIT cells migrate from the blood into the plaques.

Conditions such as obesity, chronic liver injury, and immune-mediated disorders are associated with a reduced number of circulating MAIT cells^16,9^. We also found a significant reduction in MAIT cell numbers in the blood of patients with CVD compared to healthy controls, possibly due to the observed increased cellular activation and exhaustion that may lead to apoptosis.

One of our important findings shows that human plaque MAIT cells display an activated phenotype, characterized by increased expression of activation markers CD25 and CD69, along with features of exhaustion, including elevated TIM-3 expression. However, this phenotype is not unique to MAIT cells, as many other immune cells, particularly T cells, are also highly activated within plaques^28^.

Furthermore, bulk RNA sequencing of MAIT cells sorted from both PBMCs and plaques revealed a transcriptional profile in plaque-derived MAIT cells consistent with activation and inflammation when compared to circulating counterparts from the same patients, further supporting their activated state. While the exact fate of circulating MAIT cells in chronic inflammatory conditions remains uncertain, our data support the hypothesis that MAIT cells are recruited from the bloodstream to inflamed vascular tissues, where they might contribute to local immune activation and possibly promote atherosclerosis progression.

Using mouse models under atherogenic conditions, MAIT cells accumulated in organs affected by the HC diet, particularly the liver, and were also detected within atherosclerotic plaques, albeit to a lesser extent. These observations suggest that MAIT cells may contribute to local inflammation through cytokine and chemokine production. Similarly, human atherosclerotic plaques contained a higher proportion of activated MAIT cells, which showed increased expression of chemokines compared to matched peripheral blood. Additionally, circulating MAIT cells from patients with CVD exhibited a more activated phenotype than those from healthy donors. These findings reinforce the translational relevance of our mouse model data to human pathology.

Given the systemic nature of atherosclerosis, MAIT cells may also contribute to immune activation in peripheral tissues, and this broader immune response could, in turn, influence plaque progression. Supporting this possibility, murine MAIT cells exhibited an activated phenotype in multiple organs, including the bone marrow, liver, and spleen, under atherogenic conditions.

Functionally, MAIT cells represent a unique T cell subset with innate-like features, capable of cytokine production independent of antigen specificity. In mice, these cells express a semi-invariant TCR composed of the Vα19-Jα33 TCRα chain, commonly paired with Vβ6 ^29^. To facilitate the study of MAIT cells in mice, where they are naturally scarce compared to humans, transgenic (Tg) mice expressing the invariant mVα19-Jα33 TCRα chain were initially generated^19^. Although these transgenic mice exhibit a high level of MAIT cells, they also display altered T cell ontogeny, introducing an MR1-dependent bias in TCRβ chain usage toward Vβ6 and Vβ8^19^. To overcome this limitation, the same group developed B6-MAIT^CAST^ mice, which harbor a unique CAST genetic locus that promotes a natural increase in MAIT cell numbers without perturbing T cell development or introducing ontogenic bias^11^.

The role of MAIT cells in atherosclerosis remains poorly understood. A recent study using Tg Vα19 mice suggested that MAIT cells exert an atheroprotective effect by promoting cholesterol clearance^20^. However, this transgenic model not only increased MAIT cell numbers but also potentially affected MAIT cell development and function in an MR1-dependent manner^19^. The non-specific potential impact of Vα19 transgene and the lack of an appropriate negative control group (e.g., Vα19^+/−^ MR1^−/−^ mice), limits the ability to draw definitive conclusions about the role of MAIT cells in atherosclerosis. In contrast, our study employed B6-MAIT^CAST^ mice, which naturally have increased MAIT cell numbers, alongside B6-MAIT^CAST^ MR1^−/−^ mice as proper MAIT cell-deficient negative controls. We found that the naturally elevated MAIT cell frequency in Ldlr^−/−CAST^ mice was associated with increased atherosclerosis, particularly in the thoracic aorta, compared to both Ldlr^−/−CAST^ MR1^−/−^ mice, which lack MAIT cells, and Ldlr^−/−^ littermates, which exhibit low MAIT cell levels. This effect was more prominent in female mice than in males, which may be due higher MAIT cell activation in females, as has also been reported in other settings, such as infection^30^. This likely reflects the influence of sex hormone on immune activation dynamics. Notably, this pro-atherogenic effect occurred without significant changes in plasma cholesterol levels, suggesting that inflammation, rather than lipid metabolism, drives the contribution of MAIT cells to atherosclerosis.

MAIT cells may also contribute to complications associated with atherosclerosis such as liver fibrosis and MI. One systemic effect of MAIT cells under atherogenic conditions is their observed impact on fibrosis within the liver, where they have been previously shown to play a deleterious role^16,22^. Consistently, the lack of MAIT cells under atherogenic conditions was associated with reduced atherosclerosis, decreased liver fibrosis, improved cardiac ejection fraction, and smaller infarct size following induction of MI. These results indicate that MAIT cells exert deleterious effects, influencing not only atherosclerosis but also its associated complications. This may also reflect broader MAIT cell-mediated systemic inflammatory responses that contribute to both atherosclerosis and related comorbidities.

To dissect the underlying immune mechanisms, we analyzed immune cell populations and found that MAIT cell levels are associated with those of monocytes/macrophages, which are key drivers of inflammation in atherosclerosis^31^, and have also been shown to interact with MAIT cells to promote liver fibrosis^16,22^. Interestingly, positive correlations were observed between circulating MAIT cells and monocyte numbers in both human blood and plaques, suggesting a potential link between these cell populations. Among the various factors produced by MAIT cells that can affect monocyte/macrophage survival and recruitment are hematopoietic growth factors such as GM-CSF and M-CSF, as well as several chemokines^9,32–35^. Notably, human plaque-resident MAIT cells also showed enriched functional pathways related to activation and chemotaxis, as indicated by increased expression of M-CSF and chemokine-related genes, including CCL3, CCL4, and CCL20, compared to blood-derived MAIT cells. These observations suggest that MAIT cells may contribute to the recruitment and local modulation of monocytes and macrophages within atherosclerotic plaques. Consistent with this, mouse models with elevated MAIT-cell numbers displayed increased circulating monocytes, whereas models with reduced or absent MAIT cells showed a blunted rise in circulating monocytes and decreased macrophage accumulation within plaques. In line with these findings, monocyte populations and their progenitors in the bone marrow were also altered. Within this compartment, MAIT cells identified using 5-OP-RU–loaded MR1 tetramers displayed an activated phenotype, characterized by elevated CD69 and CD25 expression under HC conditions. Moreover, bulk RNA-sequencing of bone-marrow-sorted MAIT cells under atherogenic conditions further revealed upregulation of gene programs associated with immune-cell activation and the positive regulation of immune responses. These transcriptional signatures, related to myelopoiesis and inflammation, resemble chronic activation within the bone-marrow niche, potentially skewing HSC differentiation toward myeloid lineages^36^. This shift is consistent with the decreased abundance of myeloid progenitors observed in MAIT-deficient mice under HC diet, revealing a novel function of MAIT cells in the bone marrow that may contribute to monocytosis under atherogenic conditions. Taken together, these observations suggest a close relationship between MAIT cells and monocyte/macrophage lineage, consistent with observations in other inflammatory diseases such as liver fibrosis^21^.

To further demonstrate the contribution of monocytes/macrophages to MAIT cell-mediated pro-atherogenic effects, we used the CCR2^−/−^ model, which impairs monocyte egress from the bone marrow, resulting in reduced circulating monocyte levels and abolishing the differences in plaque size between Ldlr^−/−CAST^ MR1^−/−^ mice and Ldlr^−/−CAST^ MR1^+/+^ mice. Under CCR2^+/+^ conditions, Ldlr^−/−CAST^ MR1^−/−^ mice displayed significantly smaller plaque size compared to Ldlr^−/−CAST^ MR1^+/+^ mice. These findings suggest that the pro-atherogenic role of MAIT cells is mediated, at least in part, through monocytes/macrophages.

In conclusion, our study demonstrates that MAIT cells are enriched among CD3⁺ T cells within human atherosclerotic plaques and exhibit an activated phenotype. Using mouse models, we show that high levels of MAIT cells have pro-atherogenic effects. Mechanistically, these effects appear to be mediated through altered monocyte and macrophage dynamics, highlighting a key role for MAIT cells in promoting vascular inflammation and atherogenesis. Conversely, their absence leads to a significant reduction in atherosclerosis and its associated complications, suggesting that targeting this cell type could help reduce inflammation and chronic inflammatory diseases such as atherosclerosis.

## Materials and Methods

### Human samples

Human samples were collected from patients undergoing carotid endarterectomy (CEA) (Patients’ clinical data are presented in the **Supp. Table 1**) after informed written consent was obtained with the approval of the “Comité de Protection des Personnes Ile-de-France II” (registration number 2016-13-09 MS2). Patients were excluded if they had a current infection, autoimmune disease, active or recurrent cancer, were on dialysis, had evidence of recent gastrointestinal bleeding, were treated with immunosuppressive drugs in the past 30 days, or had a human immunodeficiency virus (HIV) infection. Blood from healthy volunteers was obtained through a formalized agreement with French Blood Agency (Etablissement Français du Sang, agreement number CCPSL 2024-2027-001). Data collection and analyses were performed anonymously.

Peripheral white blood cell counts were obtained for each patient using an electrical impedance–based hematology analyzer. Monocyte counts measured the day before CEA were retrieved from the clinical laboratory records.

### Isolation of human peripheral blood mononuclear cells (PBMCs)

Blood was drawn into precoated EDTA tubes. Within two hours after sample collection, PBMCs were obtained by layering whole human blood onto Histopaque-1077 (10771, Sigma-Aldrich) and centrifuging for 30 min at 400 × g at room temperature (RT) using the lowest acceleration and brake settings, according to the manufacturer’s instructions. Subsequently, samples were washed twice in PBS (phosphate-buffered saline) and resuspended in RPMI (Roswell Park Memorial Institute medium) supplemented with 10% FCS (fetal calf serum).

### Leukocyte isolation from human CEA samples

Leukocyte isolation from human plaques was performed as previously described^37^. In particular, CEA samples were placed in RPMI supplemented with 10% FCS and stored on ice. To obtain a single-cell suspension, samples were minced using scissors and incubated for 60 min at 37 °C in pre-warmed RPMI containing 10% FCS, 0.3 mg/ml Collagenase type II (C6885, Sigma-Aldrich), 1 mg/ml Collagenase type IV (C5138, Sigma-Aldrich), 0.3 mg/ml Collagenase type XI (C7657, Sigma-Aldrich), 0.3 mg/ml Hyaluronidase (H3506, Sigma-Aldrich), and 0.3 mg/ml DNase I (10104159001, Roche). Next, cells were filtered through a 70 µm cell strainer, followed by a 40/70% Percoll (GE17-0891-02, Cytiva) density gradient centrifugation (20 min at 800 × g at RT, lowest acceleration, and no brake) to enrich leukocytes. Cells were washed and resuspended in RPMI + 10% FCS.

### Flow cytometry analysis of human samples

Following leukocyte isolation, cells were washed with PBS and incubated with FVD eFluor 506 for 30 minutes at 4 °C to exclude dead cells, followed by incubation with Fc Block to prevent nonspecific antibody binding. For stimulation, 3 × 10⁶ cells per sample were seeded into a 96-well plate and incubated for 4 hours at 37 °C with 5% CO₂ in RPMI medium supplemented with 10% FCS and Leukocyte Activation Cocktail with BD GolgiPlug (550583, BD), according to the manufacturer’s instructions. After stimulation, cells were washed and processed for flow cytometry staining. For surface staining, an antibody master mix was added directly on top of the Fc Block, and cells were incubated for 30 minutes at 4 °C. For MAIT cell staining, the tetramer loaded with 5-OP-RU or 6-FP as negative control was incubated separately from the other surface antibodies for 40 min. For cytoplasmic staining (IFN-γ, Granzyme B, and IL-17), cells were fixed in IC Fixation Buffer (00-8222, Thermo Fisher) for 30 minutes at RT. All antibodies were titrated to determine the optimal concentration and minimize signal spreading. For intracellular staining of IFN-γ, Granzyme B, and IL-17, unstimulated cells were included as negative controls to establish gating thresholds.

Absolute cell counts were obtained by adding 10 µl of Precision Count Beads (424902, BioLegend) to each sample and calculated with the following formula: Number of cells = number of acquired cells/number of acquired beads x beads/well x dilution factor. Data was acquired on an LSRFortessa flow cytometer (BD) and analyzed with FlowJo version 10.10.0 (BD).

#### Anti-human antibodies

**Anti-human antibodies:**
| Fluorophore | Antigen | Clone | Concentration | Vendor | Catalog No. |
| --- | --- | --- | --- | --- | --- |
| NA | Fc Block | Fc1 | 10 μg/ml | BD | 564220 |
| FVD eFluor 506 | live/dead | NA | 1:1000 | Thermo Fisher | 65-0866 |
| FITC | bcl-2 | Bcl-2/100 | 1:40 | BD | 340575 |
| KIRAVIA blue 520 | CD69 | FN50 | 1.25 μg/ml | BioLegend | 310956 |
| PE | MR1 5-OP-RU | NA | 1.25 μg/ml | NIH Tetramer Core Facility | NA |
| PE | 6-FP | NA | 1.25 μg/ml | NIH Tetramer Core Facility | NA |
| PerCP-Cy5.5 | CD8 | SK1 | 1.25 μg/ml | BioLegend | 344710 |
| PE-Cy7 | Granzyme B | QA16A02 | 0.625 μg/ml | BioLegend | 372214 |
| PE-Cy7 | PD1 | EH12.2H7 | 5 μg/ml | BioLegend | 329918 |
| APC | TCR Va7.2 | 3C10 | 2.5 μg/ml | BioLegend | 351708 |
| Alexa Fluor 700 | CD4 | OKT4 | 0.625 μg/ml | BioLegend | 317426 |
| APC/Fire 750 | CD3 | SK7 | 2.5 μg/ml | BioLegend | 344840 |
| BV421 | IL-17A | BL168 | 0.3125 μg/ml | BioLegend | 512322 |
| BV421 | Tim-3 | F38-2E2 | 5 μg/ml | BioLegend | 345008 |
| BV510 | CD14 | M5E2 | 2.5 μg/ml | BioLegend | 301842 |
| BV510 | CD19 | HIB19 | 2.5 μg/ml | BioLegend | 302242 |
| BV605 | IFN-γ | 4S.B3 | 1.25 μg/ml | BioLegend | 502536 |
| BV605 | CD25 | BC96 | 5 μg/ml | BioLegend | 302632 |

### Mice

All experiments were conducted in accordance with the ethical guidelines for animal experimentation set by the University of Paris Cité (CEEA 34) and the National Charter on the Ethics of Animal Experimentation from the French Ministry of Higher Education and Research (reference MESR no. 01373.01). Both male and female mice were used in this study. Ldlr^−/−CAST^ and Ldlr^−/−CAST^ MR1^−/−^ mice were generated by crossing Ldlr^−/−^ mice (JAX:002207) with B6-MAIT^CAST^ or B6-MAIT^CAST^ MR1^−/−^ mice bred in-house; the B6-MAIT^CAST^ and B6-MAIT^CAST^ MR1^−/−^ mouse lines were kindly provided by Pr. Olivier Lantz. All mice were on a C57Bl/6 background and had been backcrossed for more than 10 generations. In general, mice (n = 5 per cage) were separated by genotype at weaning and maintained until sacrifice. Animals were housed under a 12-hour light/dark cycle with ad libitum access to water and a standard chow diet. Littermate mice (Ldlr^−/−^, Ldlr^−/−CAST^, or Ldlr^−/−CAST^ MR1^−/−^) were either kept on a normal chow diet (CD; A03, SAFE, France) or placed on a high-cholesterol (HC) diet containing 1.25% cholesterol (E15106-347, SSNIFF, Germany), starting at 7 weeks of age for up to 14 weeks.

To generate chimeric mice, 10-week-old Ldlr^−/−^ mice were subjected to medullar aplasia by lethal whole-body irradiation (9.5 Gray) and transplanted with bone marrow cells isolated from femurs and tibias of sex and age-matched either B6 or B6-MAIT^CAST^ mice. In another set of experiments 10-week-old C57bl/6J Ldlr^−/−CAST^ and Ldlr^−/−CAST^ MR1^−/−^ mice, were subjected to medullar aplasia by lethal whole-body irradiation (9.5 Gray). The mice were repopulated with an intravenous injection of bone marrow cells isolated from femurs and tibias of sex and age-matched CCR2^+/+^ or CCR2^−/−^. After 4 weeks of recovery, mice were fed a HC diet.

In some experiments, acute myocardial infarction was induced in mice by permanent ligation of the left anterior descending coronary artery. Cardiac function and remodeling were analyzed by using echocardiography and immunohistochemistry ^38^. Briefly, mice were anesthetized using Ketamine (100 mg/kg body weight) and Xylazine (10 mg/kg body weight) via intra-peritoneal injection, intubated and ventilated using a small-animal respirator. The chest wall was shaved and a thoracotomy was performed in the fourth left intercostal space. The pericardial sac was removed, and the left anterior descending artery was permanently ligated using a 7/0 monofilament suture (Peters surgical, France) at the site of its emergence close to the left atrium. The thoracotomy was closed with 6/0 monofilament sutures. The endotracheal tube was removed once spontaneous respiration was resumed, and mice were placed on a warm pad at 37 °C until awakened. All experiments were performed with 10-week-old male animals and were carried out using age and littermate matched groups without randomization.

All animal procedures were performed in accordance with the European Directive 2010/63/EU and institutional guidelines approved by the local ethics committee (’Comité d’Éthique en Expérimentation Animale’) under protocol number APAFIS #33149-2021073015348753.

### Echocardiography

Transthoracic echocardiography in mice was performed using a Vevo 2100 imaging system (VisualSonics, Fujifilm) equipped with a high-frequency transducer (MS400, VisualSonics, Fujifilm). The imaging platform and ultrasound gel were pre-heated to 40 °C. Anesthesia was induced with isoflurane at a concentration of 1–2 vol% in oxygen at a flow rate of 1 L/min. Once stable sedation was achieved, mice were positioned with all four limbs secured to ECG electrodes to allow continuous heart rate monitoring, and chemical hair removal was performed at the imaging site. Core body temperature was monitored using a rectal thermometer. Anesthesia was adjusted throughout the procedure to maintain a heart rate of 500–600 beats per minute and a body temperature between 36.5 °C and 38.0 °C. Echocardiographic recordings were acquired in parasternal long- and short-axis views. Image analysis and calculations were performed using dedicated software (Vevo LAB, VisualSonics, Fujifilm). The investigator was blinded to group assignment. Percentages of ejection fraction (%EF) were calculated^39^.

### Plasma cholesterol measurement

Plasma was obtained from EDTA blood samples after centrifugation for 10 min at 2000 g at 4°C. Cholesterol levels were measured by using a commercially available colorimetric test according to the manufacturer’s instructions (1 1300 99 10 023, Cholesterol FS, DiaSys).

### Aorta immunofluorescence whole mount staining

Aortas previously stained with Oil Red O and mounted in Fluoromount were rehydrated overnight in 1X PBS to remove the coverslip, then encircled with a hydrophobic barrier. Samples were washed in 1X PBS, permeabilized, and blocked for 1 hour in 1X PBS containing 3% BSA and 0.3% Triton X-100 before incubation with the primary antibody (rat anti-MOMA-2, Sigma-Aldrich #MAB1852, 1:100, 48 h, 4 °C). After washing with 1X PBS containing 0.3% Triton X-100, tissues were incubated with the fluorescent secondary antibody (donkey anti-rat Cy5, Jackson ImmunoResearch #712-175-153, 48 h, 4 °C), followed by DAPI staining. Aortas were then washed and mounted in Fluoromount. Imaging was performed using an inverted fixed-stage TCS SP8 confocal microscope (Leica Microsystems, France) equipped with 405, 561, and 640 nm laser lines. Images were acquired with a 20X objective and captured using a hybrid detector in photon-counting z-stack mode. MOMA-2-positive macrophage areas were quantified using QuPath 4.0.

### Evaluation of liver fibrosis

Sirius Red staining was performed on 4-µm-thick formalin-fixed paraffin-embedded tissue sections. Sirius Red-stained areas from ten fields (magnification 10×) from each mouse were quantified with ImageJ.

### Evaluation of infarct size

For evaluation of cardiac remodeling, hearts were excised, rinsed in PBS snap-frozen in isopentane chilled with liquid nitrogen. Hearts were cut by a cryostat (CM 3050S, Leica) into 10μm thick sections of the entire heart from apex to root. Masson’s trichrome staining was performed for infarct size evaluation. Infarct size was calculated as the ratio of total infarct circumference divided by total left ventricle circumference.

### Murine tissue processing and flow cytometry

After perfusion of the mouse with 10 ml of ice-cold PBS, aortas were minced with scissors and incubated in prewarmed RPMI containing 450 U/ml Collagenase I (C0130, Sigma-Aldrich), 125 U/ml Collagenase XI (C7657, Sigma-Aldrich), 60 U/ml Hyaluronidase (H3506, Sigma-Aldrich), and 60 U/ml DNAse I (10104159001, Roche) for 60 minutes at 37 °C. The digested tissue was then washed and filtered through a 70 µm cell strainer. Blood leukocytes were obtained by treating whole blood twice with 1×RBC lysis buffer (00-4300, Thermo Fisher) for 5 minutes and 3 minutes at room temperature (RT), respectively. Bone marrow was flushed from the left femur with a 26G needle. Bone marrows, livers and spleens were mashed through a 70 µm cell strainer. Liver suspensions were diluted in 10 ml of 42% Percoll solution (GE17-0891-02, Cytiva) and centrifuged for 20 minutes at 800 g at RT (with lowest acceleration and no brake). Bone marrow, liver and spleen cells were then incubated in RBC lysis buffer for 3 minutes at RT. After washing, cells were processed for flow cytometry staining. Therefore, cells were incubated with FVD eFluor 506 for 30 minutes at 4 °C to exclude dead cells. For surface staining, an antibody master mix was added directly on top of the Fc Block, and cells were incubated for 30 minutes at 4 °C. For bone marrow progenitor staining, Fc blocking was not performed because it would compete with the anti-CD16/32 antibody included in the master mix. For MAIT cell staining, the tetramer was incubated simultaneously with other surface antibodies for 40 minutes at RT. For nuclear staining (e.g., RORγt, FOXP3, T-bet), cells were permeabilized using Fixation/Permeabilization Buffer (00-5523, Thermo Fisher), followed by staining in Permeabilization Buffer (00-8333, Thermo Fisher) for 30 minutes at RT. All samples were kept on ice until further processing. Absolute cell counts were obtained by adding 10 µl of Precision Count Beads (424902, BioLegend) to each sample and calculated using the formula: Number of cells = (number of acquired cells / number of acquired beads) × beads per well × dilution factor. Data were acquired on an LSRFortessa flow cytometer (BD) and analyzed using FlowJo version 10.10.0 (BD).

#### Anti-mouse antibodies

**Anti-mouse antibodies:**
| **Fluorophore** | **Antigen** | **Clone** | **Concentration** | **Vendor** | **Catalog No.** |
| --- | --- | --- | --- | --- | --- |
| NA | Fc Block | 2.4G2 | 10 µg/ml | BD | 553142 |
| FVD eFluor 506 | live/dead | NA | 1:1000 | Thermo Fisher | 65-0866 |
| FITC | CD45 | 30-F11 | 2.5 µg/ml | BioLegend | 103108 |
| Alexa Fluor 488 | CD11b | M1/70 | 5 µg/ml | BioLegend | 101219 |
| KIRAVIA blue 520 | CD69 | H1.2F3 | 1.25 µg/ml | BioLegend | 104554 |
| PE | Ly-6G | 1A8 | 2.5 µg/ml | BioLegend | 127608 |
| PE | MR1 5-OP-RU | NA | 2.5 µg/ml | NIH Tetramer Core Facility | NA |
| PE | 6-FP | NA | 2.5 µg/ml | NIH Tetramer Core Facility | NA |
| PE | CD48 | HM48-1 | 2 µg/ml | BioLegend | 103406 |
| PE | c-Kit | 2B8 | 2 µg/ml | BioLegend | 105807 |
| PerCP-Cy5.5 | CD19 | 6D5 | 0.625 µg/ml | BioLegend | 115534 |
| PerCP-Cy5.5 | CD8a | 53-6.7 | 0.3125 µg/ml | BioLegend | 100734 |
| RB780 | T-bet | O4-46 | 1.25 µg/ml | BD | 569089 |
| PE-Cy7 | CD3ε | 145-2C11 | 2.5 µg/ml | BioLegend | 100320 |
| PE-Cy7 | Granzyme B | QA16A02 | 0.625 µg/ml | BioLegend | 372214 |
| PE-Cy7 | PD-1 | RMP1-30 | 0.3125 µg/ml | BioLegend | 109110 |
| PE-Cy7 | CD45R | RA3-6B2 | 2 µg/ml | BioLegend | 103221 |
| PE-Cy7 | Ter-119 | TER-119 | 2 µg/ml | BioLegend | 116221 |
| PE-Cy7 | Gr1 | RB6-8C5 | 2 µg/ml | BioLegend | 108415 |
| APC | CD64 | X54-5/7.1 | 2.5 µg/ml | BioLegend | 139306 |
| APC | CD115 | AFS98 | 0.3125 µg/ml | BioLegend | 135510 |
| APC | RORγt | B2D | 1.25 µg/ml | Thermo Fisher | 17-6981-82 |
| APC | CD135 | A2F10.1 | 2 µg/ml | BD | 560718 |
| APC | CD44 | IM7 | 1.25 µg/ml | BioLegend | 103012 |
| Alexa Fluor 647 | CD34 | RAM34 | 2 µg/ml | BD | 560233 |
| Alexa Fluor 700 | CD4 | RM4-5 | 0.625 µg/ml | BioLegend | 100536 |
| Alexa Fluor 700 | CD45 | 30-F11 | 5 µg/ml | BioLegend | 103128 |
| APC-Cy7 | c-Kit | 2B8 | 2 µg/ml | BioLegend | 105826 |
| APC/Fire 750 | Ly-6C | HK1.4 | 2.5 µg/ml | BioLegend | 128046 |
| APC/Fire 750 | CD3ε | 145-2C11 | 2.5 µg/ml | BioLegend | 100362 |
| APC/Fire750 | Sca1 | D7 | 2 µg/ml | BioLegend | 108145 |
| BV421 | CD11b | M1/70 | 0.3125 µg/ml | BioLegend | 101236 |
| BV421 | TCR β | H57-597 | 1.25 µg/ml | BioLegend | 109230 |
| BV421 | CD16/32 | 2.4G2 | 2 µg/ml | BD (OptiBuild) | 752948 |
| V450 | Lineage Antibody Cocktail | 500A2; M1/70; RA3-6B2; TER-119; RB6-8C5 | 1:100 | BD | 561301 |
| BV510 | CD11b | M1/70 | 0.625 µg/ml | BioLegend | 101263 |
| BV510 | CD11c | N418 | 0.625 µg/ml | BioLegend | 117338 |
| BV510 | CD19 | 6D5 | 1.25 µg/ml | BioLegend | 115546 |
| BV510 | B220 | RA3-6B2 | 2.5 µg/ml | BioLegend | 103248 |
| BV605 | CD8a | 5H10-1 | 1.25 µg/ml | BioLegend | 100802 |
| BV605 | CD25 | PC61 | 0.625 µg/ml | BioLegend | 102036 |
| BV711 | CD150 | TC15-12F12.2 | 2 µg/ml | BioLegend | 115941 |
| BUV395 | Sca1 | E13-161.7 | 2 µg/ml | BD | 744328 |

### Murine tissue processing and analysis for plaque histology

Mice were perfused with 10 ml of ice-cold PBS to clear the blood from the circulation. Aortic roots were embedded in OCT compound and snap-frozen in isopentane chilled with liquid nitrogen. Serial 10 µm-thick cryosections of the aortic root were obtained using a cryotome and used for subsequent staining. Sections were stained with Oil Red O (O0625, Sigma-Aldrich) or Picro-Sirius Red (ab246832, Abcam). For immunohistochemistry, sections were stained with either anti-CD3 antibody (polyclonal, A0452, Agilent, 1:200) or anti-macrophage/monocyte antibody (clone MOMA-2, MAB1852, Merck, 1:100), followed by appropriate secondary antibodies: Alexa Fluor 594 Goat Anti-Rat IgG (H+L) (112-585-167, 1:100) or Alexa Fluor 647 Goat Anti-Rabbit IgG (H+L) (111-605-144, 1:200; both from Jackson ImmunoResearch). Quantification within atherosclerotic lesions was performed in cross-sectional areas throughout the whole aortic root, which represents ∼6-8 sections per mouse, and appropriate negative controls were used.

Image analysis was performed in a blinded manner using QuPath^40^. For en face analysis, aortas were fixed in 10% formalin, cleaned, cut longitudinally, and stained with Oil Red O. Images were acquired and analyzed using Fiji software^41^.

### MAIT cell sorting & bulk RNA sequencing

MAIT cells were sorted using a FACSAria II cell sorter equipped with a 70-µm nozzle (BD) from plaque samples obtained from four patients. We also sorted MAIT cells from bone marrow using MR1 tetramers from five mice fed either a CD or HC diet for 1 week.

The post-sort purity was analyzed by re-running a sorted sample after the addition of a viability dye and was 98.97 %. Between 160 and 1000 cells were sorted using fluorescence-activated cell sorting (FACS) and deposited into 0.2 mL tubes containing lysis buffer supplemented with 2U of Protector RNase Inhibitor and oligo(dT) primer for reverse transcription initiation, utilizing Smart technology. The resulting cDNA was then amplified for 17 cycles. cDNA shape was controlled using an HS DNA chip (Bioanalyzer 2100 / Agilent) to ensure a medium size around 1000 nucleotides, indicating high RNA quality input.

Quality control and quantification of the cDNA samples were conducted using a Qubit Fluorometer (Thermo Fisher Scientific. Q33327) and an HS DNA chip on a Bioanalyzer 2100 (Agilent). All samples passed the quality control step. Libraries were prepared starting with 1 ng of cDNA using the Nextera XT DNA Library Prep Kit (Illumina). Subsequent quality control and quantification of the libraries were performed using a Qubit Fluorometer (Thermo Fisher Scientific. Q33327) and an HS DNA chip on a Bioanalyzer 2100 (Agilent). The libraries were sequenced on an Illumina NextSeq 500 instrument using 75-base pair read lengths with V2 chemistry in a paired-end mode.

Following sequencing. primary data analysis involved demultiplexing and quality control of raw data using AOZAN software (ENS. Paris) based on FastQC modules (version 0.11.5). The resulting FASTQ files were aligned using the STAR algorithm (version 2.5.2b). and alignment quality was assessed using Picard tools (version 2.8.1). Reads were counted using FeatureCounts (version Rsubread 1.24.1) and statistical analyses on read counts were performed using the DESeq2 package (version 1.14.1) to identify differentially expressed genes between experimental conditions All raw human RNA sequencing data files have been deposited in the Gene Expression Omnibus (GEO) database under accession number GSE300578. Raw mouse RNA sequencing data files have been deposited in the GEO database under accession number GSE310744.

### Bulk RNA sequencing analysis

Raw read counts were normalized using the DESeq2 package. Principal component analysis was performed on the 1000 most variable genes. Differential expression analyses were conducted with DESeq2 to compare whole blood from patients versus healthy controls (1309 genes with adjusted p-value < 0.05) and plaque tissue versus whole blood from the same patients (1088 genes with adjusted p-value < 0.05). For each comparison, the top 50 genes with the lowest adjusted p-values were used to generate heatmaps. Genes with an adjusted p-value < 0.001 were subjected to GO biological process enrichment analysis, using the top 20 terms ranked by adjusted p-value.

For mouse data, genes identified as differentially expressed by DESeq2 with a nominal p-value < 0.02 were subjected to functional enrichment analysis using Metascape (www.metascape.org). Over-represented Gene Ontology Biological Processes and pathway categories were computed using default settings, including hierarchical clustering and enrichment score calculation. Normalized expression values were obtained from variance-stabilizing transformation (VST) of DESeq2-processed count data. A GM-CSF–related gene signature was quantified using two complementary approaches: (i) computation of a mean z-score across all signature genes per sample, and (ii) single-sample gene set enrichment analysis (ssGSEA) using the GSVA framework. For each method, scores were compared between control and HC groups using Kruskal–Wallis tests.

### Statistical analysis

Statistical analysis was performed using GraphPad Prism version 9.5.0. Outliers were identified by utilizing the ROUT method with Q = 1 %. For the comparison of 2 groups, the Mann-Whitney test was used, and for paired analysis, the Wilcoxon test was used. For more than two groups, the Kruskal-Wallis test followed by Dunn’s multiple comparisons was used. Correlations were calculated using the Spearman correlation. Graphs show the mean ± SEM. Two-sided p-value < 0.05 was considered significant.

## Acknowledgements

This work was supported by Inserm, Agence Nationale de la Recherche (ANR-21CE14-0007-01 to S.T., S.L., and A.T.), Federation Française de Cardiologie (FFC) (to S.T.), and Fondation De France (FDF) (to S.T.), and Fondation pour la Recherche Médicale (FRM) (to H.A.O. and S.T.). T.R. received a scholarship from DFG (Walter Benjamin program, project number 49093744). T.N. was the recipient of FRM scholarship for his thesis. N. M. and M.C. were the recipients of a scholarship from FDF for their thesis. M.C. was the recipient of a scholarship from Nouvelle Societé Française d’Athérosclerose (NSFA) for the 4th year of her thesis. N.M. is the recipient of a scholarship from FRM for the 4th year of her thesis. We thank members of our animal and histology Facilities. We thank the Genomics Core Facility of the Institut Cochin for NGS data production and analysis.

## Author contributions statement

T.R. contributed to the experimental design, performed most of the experiments, analyzed the data, and prepared the figures. T.N., N.M., E.B., M.C., P.L., C.H., K.D., M.L. and R.A. assisted with selected experiments. C.K. contributed to the MAIT cell sorting experiments. F.L. supervised sample processing and bulk RNA sequencing experiments. M.D. performed the statistical analysis of the bulk RNA sequencing data. L.W. and T.M. provided human samples and clinical information, and contributed to the interpretation of human data. P.J., J.M.A, S.E.B. performed surgery. O.S., C.C., M.P., A.T., H.A.O., and Z.M. provided input and discussed the results. O.L. shared his expertise on MAIT cells, provided the B6-MAIT^CAST^ and B6-MAIT^CAST^ MR1^−/−^ mice, and participated in data interpretation and discussion. P.A. and J.S.S. performed and analyzed the myocardial infarction experiments, contributed to result interpretation, and participated in discussions. C.C. and S.L. carried out liver assessments, analyzed the data, and contributed to interpretation and discussion. S.T. conceived and designed the study, analyzed and interpreted the data, and wrote the manuscript.

## Competing interest statement

The authors declare no competing interests.

## Notes

### Competing Interest Statement

The authors have declared no competing interest.

