## Supplemental material for "Mucosal-Associated Invariant T Cells Promote Atherosclerosis Through Monocyte-Driven Inflammation"

### Supplementary Figure legends

**Supplementary Fig. 1 Characterization of human MAIT cells by flow cytometry. A**. Example of the MAIT cell gating strategy in humans. **B.** Schematic representation of the experimental design; CEA (carotid endarterectomy). **C.** Correlation of MAIT cells in the blood and plaque (n = 13). **D-F.** Representative pseudocolor plots and quantification of % of granzyme $(GrzB)^+$ MAIT cells, IFN-$\gamma^+$ MAIT cells, and $IL17^+$ MAIT cells in blood and plaque from the same patients (n = 13). **G.** Representative pseudocolor plots and quantification of $CD8^+$ and double negative (DN) $CD4^-CD8^-$ MAIT cells in the blood from CTRLs (n = 15) and patients (n = 14). **H.** Correlations of blood MAIT cell numbers with gMFI of $CD25^+$, $CD69^+$, and $TIM3^+$ MAIT cells. **I.** Correlations of MAIT cells in the blood and plaque with monocyte counts from the same patient (n = 14). gMFI, geometric mean fluorescence intensity. Individual data are presented as before-after or scatter plots, with the mean and SEM. The p-values were determined using the Wilcoxon test for data presented in a before-after plot or the Mann-Whitney test for data presented in a scatter plot. Correlations were calculated using Spearman correlation. Correlation graphs show the 95% confidence interval in grey.

**Supplementary Fig. 2. Characterization of murine MAIT cells in different tissues by flow cytometry. A.** Example of the MAIT cell gating strategy in mice. **B.** Representative pseudocolor plots and quantification of blood MAIT cell staining with 5-OP-RU tetramer in male $Ldlr^{-/-CAST}$ mice fed a chow diet (CD) or a high-cholesterol diet (HC) for eight weeks (n = 5 per group, replicated in two independent experiments). **C.** Representative pseudocolor plots and quantification of liver MAIT cell staining with 5-OP-RU tetramer in male $Ldlr^{-/-CAST}$ mice fed a CD or a HC diet for one week (n = 5 per group, replicated in two independent experiments). **D.** Representative pseudocolor plots and quantification of $CD4^+$, $CD8^+$, and double-negative (DN; $CD4^-CD8^-$) MAIT cells from the spleen of male $Ldlr^{-/-CAST}$ mice fed a CD or a HC diet for eight weeks (n = 10 per group, pool of two independent experiments). **E**. Pseudocolor plots and quantification of $CD4^+$, $CD8^+$, and DN MAIT cells from aortas of male $Ldlr^{-/-CAST}$ mice fed a CD or a HC diet for eight weeks (n = 5 per group, replicated in two independent experiments). Individual data are presented as scatter plots, with the mean and SEM. P-values were determined using the Mann–Whitney test.

**Supplementary Fig. 3. MAIT cell activation and polarization under an atherogenic diet. A.** Representative Pseudocolor plots of FMO controls for CD69 and PD1. **B.** Representative pseudocolor plots and quantification of $CD69^+$ MAIT cells from livers of female $Ldlr^{-/-CAST}$ mice fed a chow diet (CD) or high-cholesterol diet (HC) for one week, n = 4-5 per group. **C-D.** Representative pseudocolor plots and quantification of $CD69^+$ and $PD1^+$ MAIT cells from spleens of male $Ldlr^{-/-CAST}$ mice fed a CD or a HC diet for eight weeks (n = 5 per group, replicated in two independent experiments). **E.** Representative pseudocolor plots and quantification of $CD69^+$ MAIT cells from aortas of male $Ldlr^{-/-}$

$^{CAST}$ mice fed a CD or a HC diet for eight weeks (n = 5 per group). **F.** Representative pseudocolor plots and quantification of T-bet$^{+}$ (MAIT1) and RORγt$^{+}$ (MAIT17) MAIT cells from aortas of male Ldlr$^{-/-CAST}$ mice fed a CD or a HC diet for eight weeks (n = 5 per group). **G-H.** Representative pseudocolor plots and quantification of T-bet$^{+}$ (MAIT1) and RORγt$^{+}$ (MAIT17) MAIT cells from spleens and livers of male Ldlr$^{-/-CAST}$ mice fed a CD or a HC diet for eight weeks (n = 5 per group). Individual data are presented as scatter plots, with the mean and SEM. P-values were determined using the Mann–Whitney test.

**Supplementary Fig. 4. MAIT cells play a pro-atherogenic role in mice. A.** Representative pseudocolor plots and quantification of MAIT cell staining from the liver of female Ldlr$^{-/-}$ mice reconstituted with bone marrow from female B6 or B6-MAIT$^{CAST}$ mice. **B.** Plasma cholesterol levels in irradiated female Ldlr$^{-/-}$ mice reconstituted with bone marrow from female B6 or B6-MAIT$^{CAST}$ mice, followed by a ten-week high-cholesterol (HC) diet (n=9-10 per group). **C.** Correlation of plasma cholesterol with plaque area in the thoracic aorta and aortic root. **D.** Plasma cholesterol levels in male Ldlr$^{-/-}$, Ldlr$^{-/-CAST}$, and Ldlr$^{-/-CAST}$ MR1$^{-/-}$ mice fed a HC diet for eight weeks. **E.** Representative pseudocolor plots and quantification of MAIT cell staining with 5-OP-RU tetramer in spleens of male Ldlr$^{-/-}$, Ldlr$^{-/-CAST}$, and Ldlr$^{-/-CAST}$ MR1$^{-/-}$ mice fed a HC diet for eight weeks. **F.** Brightfield images of Oil Red O-stained thoracic aortas (scale bar: 2 mm) and quantification from male Ldlr$^{-/-}$, Ldlr$^{-/-CAST}$, and Ldlr$^{-/-CAST}$ MR1$^{-/-}$ mice fed a HC diet for eight weeks (n = 10-15 per group). Individual data are presented as scatter plots, with the mean and SEM. For comparison of two groups, the Mann–Whitney test was used; for more than two groups, the Kruskal–Wallis test followed by Dunn's multiple comparisons was applied. Correlations were calculated using the Spearman correlation, and graphs show the 95% confidence interval in grey.

**Supplementary Fig. 5. Impact of MAIT cells on atherosclerosis size and composition in the aortic root. A.** Brightfield images and quantification of Oil Red O-stained aortic root sections from female Ldlr$^{-/-}$, Ldlr$^{-/-CAST}$, and Ldlr$^{-/-CAST}$ MR1$^{-/-}$ mice fed a high-cholesterol (HC) diet for eight weeks (scale bar: 200 µm; n = 13- 25 per group). **B.** Brightfield images and quantification of Oil Red O-stained aortic root sections from male Ldlr$^{-/-}$, Ldlr$^{-/- CAST}$, and Ldlr$^{-/-CAST}$ MR1$^{-/-}$ mice fed a HC diet for eight weeks (scale bar: 200 µm; n = 19-30 per group; pool of three independent experiments). **C.** Representative immunofluorescence images and quantification of macrophage accumulation stained with MOMA-2 in aortic root sections from female Ldlr$^{-/-}$, Ldlr$^{-/-CAST}$, and Ldlr$^{-/-CAST}$ MR1$^{-/-}$ mice fed a HC diet for eight weeks (scale bar: 200 µm; n = 13- 25 per group). **D.** Representative immunofluorescence images and quantification of T-cell accumulation stained with CD3 in aortic root sections from female Ldlr$^{-/-}$, Ldlr$^{-/-CAST}$, and Ldlr$^{-/-CAST}$ MR1$^{-/-}$ mice fed a HC diet for eight weeks (scale bar: 200 µm, n = 13- 25 per group, pool of three independent experiments). **E.** Brightfield images and quantification of Sirius Red-stained aortic root sections from female Ldlr$^{-/-}$ (n = 25), Ldlr$^{-/-CAST}$, and Ldlr$^{-/-CAST}$ MR1$^{-/-}$ mice fed a HC diet for eight weeks (scale bar: 200 µm, , n = 13-25 per group, pool of three independent experiments) **F.** Representative pseudocolor plots and quantification of spleen CD25$^{+}$ MAIT cells from male (n = 15) and female (n = 10) *Ldlr*$^{-/-CAST}$ mice fed a HC diet for eight weeks (pool of four independent experiments). Individual data are presented as scatter plots, with the mean and SEM. P-values were determined using the Kruskal–Wallis test followed by Dunn's multiple comparisons.

**Supplementary Fig. 6. Impact of MAIT cells on atherosclerosis at a prolonged time point. A.** Experimental design. **B.** Brightfield images and quantification of Oil Red O-stained thoracic aortas from female Ldlr$^{-/-CAST}$ and Ldlr$^{-/-CAST}$ MR1$^{-/-}$ mice fed a high-cholesterol (HC) diet for fourteen weeks (n = 10 per group; scale bar: 2 mm). **C.** Brightfield images and quantification of Oil Red O-stained aortic root sections from the same female groups (n = 10 per group; scale bar: 200 µm). **D.** Brightfield images and quantification of Oil Red O-stained thoracic aortas from male Ldlr$^{-/-CAST}$ and Ldlr$^{-/-CAST}$ MR1$^{-/-}$ mice fed a HC diet for fourteen weeks (n = 8-15 per group; scale bar: 2 mm). **E.** Brightfield images and quantification of Oil Red O-stained aortic root sections from male Ldlr$^{-/-CAST}$ and Ldlr$^{-/-CAST}$ MR1$^{-/-}$ mice fed a HC diet for fourteen weeks (n = 7-10 per group; scale bar: 200 µm). **F.** Experimental design; LAD, left anterior descending. **G.** Brightfield images and quantification of Oil Red O-stained thoracic aortas from male Ldlr$^{-/-CAST}$ and Ldlr$^{-/-CAST}$ MR1$^{-/-}$ mice fed a HC diet for eight weeks and subjected to LAD ligation (n = 8-10 per group; scale bar: 2 mm). **H.** Correlation between plaque size in the thoracic aorta and fibrotic area in the liver of male Ldlr$^{-/-CAST}$ and Ldlr$^{-/-CAST}$ MR1$^{-/-}$ mice receiving a HC diet for eight weeks and undergoing LAD ligation. Individual data are presented as scatter plots, with the mean and SEM. P-values were determined using the Mann–Whitney test. Correlations were calculated using the Spearman correlation, and graphs display the 95% confidence interval in grey.

**Supplementary Fig. 7. Gating strategies of presented immune cell populations. A.** Gating strategy of leukocyte subsets in the blood. **B.** Gating strategy of leukocyte subsets in the aorta. **C.** Gating strategy of TH-subsets in the spleen. The gating on single cells was the same as shown in A and B.

**Supplementary Fig. 8. Characterization of immune cell populations in different compartments. A.** Quantification of classical (Ly6C$^{high}$) and non-classical (Ly6C$^{low}$) monocytes in the blood of female Ldlr$^{-/-}$, Ldlr$^{-/-CAST}$, and Ldlr$^{-/-CAST}$ MR1$^{-/-}$ mice fed a high-cholesterol (HC) diet for eight weeks (n = 15-20 per group; pool of four independent experiments). **B.** Quantification of neutrophils (Ly6G$^{+}$), CD4$^{+}$ lymphocytes, CD8$^{+}$ lymphocytes, and B cells (CD19$^{+}$) in the blood of the same groups. **C.** Quantification of neutrophils, CD4$^{+}$ lymphocytes, CD8$^{+}$ lymphocytes, and B cells in the aortas of female Ldlr$^{-/-}$, Ldlr$^{-/-CAST}$, and Ldlr$^{-/-CAST}$ MR1$^{-/-}$ mice fed a HC diet for eight weeks (n = 5-14 per group; pool of two independent experiments). **D.** Quantification of CD4$^{+}$ lymphocytes and CD4$^{+}$ subsets defined as TH1 (T-bet$^{+}$), TH17 (RORγt$^{+}$), and Tregs (Foxp3$^{+}$) in the spleens of female Ldlr$^{-/-}$, Ldlr$^{-/-CAST}$, and Ldlr$^{-/-CAST}$ MR1$^{-/-}$ mice fed an HC diet for eight weeks (n = 15-20 per group; pool of four independent experiments).

E. Quantification of classical (Ly6C$^{high}$) and non-classical (Ly6C$^{low}$) monocytes, neutrophils and macrophages (CD11b$^{-}$ F4/80$^{+}$) in the spleen of female Ldlr$^{-/-}$ , Ldlr$^{-/-CAST}$, and Ldlr$^{-/-CAST}$ MR1$^{-/-}$ mice fed an HC diet for eight weeks (n = 10-15 per group; pool of three independent experiments). Individual data are presented as scatter plots, with the mean and SEM. P-values were determined using the Kruskal–Wallis test followed by Dunn's multiple comparisons.

**Supplementary Fig. 9. Characterization of MAIT impacts on monocytes. A.** Representative pseudocolor plots and quantification of MAIT cell staining from the bone marrow of female Ldlr$^{-/-CAST}$ MR1$^{-/-}$ and Ldlr$^{-/-CAST}$ mice fed a chow diet (CD) or a high-cholesterol (HC) diet for one week (n = 5 per group replicated in two independent experiments). **B.-C.** Representative pseudocolor plots and quantification of bone marrow (B) CD25$^{+}$ and (C) CD69$^{+}$ MAIT cells from female Ldlr$^{-/-CAST}$ mice receiving a CD or a HC diet for one week (n = 5 per group replicated in two independent experiments). **D.-E.** Representative pseudocolor plots and quantification of bone marrow (D) CD25$^{+}$ and (E) CD69$^{+}$ MAIT cells from female Ldlr$^{-/-CAST}$ mice fed a CD or a HC diet for eight weeks (n = 5 per group). **F.-G.** Bulk RNA sequencing of MAIT cells sorted from bone marrow in Ldlr$^{-/-CAST}$ mice fed a CD or a HC diet for one week (n = 5 per group). (F) ssGSEA score corresponding to the "cellular response to granulocyte macrophage colony-stimulating factor stimulus" enriched function and (G) heatmap of the corresponding genes. Individual data are presented as scatter plots, with the mean and SEM. P-values were determined using the Mann–Whitney test.

**Supplementary Fig. 10. Gating strategies of presented immune cell populations. A.** Gating strategy for myeloid progenitors LSK (Lin$^{-}$, Sca1$^{high}$, c-Kit$^{high}$), LK (Lin$^{-}$, Sca1$^{low}$, c-Kit$^{high}$), common myeloid progenitors (CMP; Lin$^{-}$ c-Kit$^{high}$ Sca1$^{low}$ CD34$^{high}$ CD16/32$^{low}$), granulocyte-macrophage progenitors (GMP; Lin$^{-}$ c-Kit$^{high}$ Sca1$^{low}$ CD34$^{high}$ CD16/32$^{high}$) and megakaryocytic-erythroid progenitors (MEP; Lin$^{-}$ c-Kit$^{high}$ Sca1$^{low}$ CD34$^{low}$ CD16/32$^{low}$) in mouse bone marrow, where the lineage (Lin) cocktail is defined as a mixture of anti-CD3ε PE-Cy7, anti-Ter-119 PE-Cy7, anti-CD45R PE-Cy7, and anti-Gr1 PE-Cy7 antibodies (see Materials and Methods). **B.** Gating strategy of LK, LSK, hematopoietic stem cells (HSC; Lin$^{-}$ c-Kit$^{high}$ Sca1$^{high}$ CD135$^{low}$ CD48$^{low}$), including short-term (ST)-HSC (CD150$^{-}$) and long-term (LT)-HSC (CD150$^{+}$) in mouse bone marrow. In this panel, the Lin cocktail used corresponds to BD reference 561301 (see Materials and Methods).

**Supplementary Fig. 11. Characterization of bone marrow progenitors. A.** Representative pseudocolor plots of LK and LSK staining and quantification from the bone marrow of female Ldlr$^{-/-CAST}$ and Ldlr$^{-/-CAST}$ MR1$^{-/-}$ mice receiving a CD (baseline, n = 5 per group). **B.** Representative pseudocolor plots of CMP / GMP / MEP staining and quantification from the bone marrow of female Ldlr$^{-/-CAST}$ and Ldlr$^{-/-CAST}$ MR1$^{-/-}$ mice receiving a CD (baseline, n = 5 per group). **C.** Representative pseudocolor plots of hematopoietic stem cells (HSC) staining and quantification from the bone marrow of female $Ldlr^{-/-CAST}$ and $Ldlr^{-/-CAST}$ $MR1^{-/-}$ mice receiving a CD (baseline) or a HC diet for one week (n = 5 per group). **D.** Representative pseudocolor plots of short-term (ST)-HSC and long-term (LT)-HSC staining and quantification from the bone marrow of female $Ldlr^{-/-CAST}$ and $Ldlr^{-/-CAST}$ $MR1^{-/-}$ mice receiving a CD (baseline) or a HC diet for one week (n = 5 per group). Individual data are presented as scatter plots, with the mean and SEM. P-values were determined using the Mann-Whitney test. Dashed lines indicate that the experiments were not done at the same time and the Mann-Whitney test was used for comparison.

**Supplementary Fig. 12. MAIT cells impact atherosclerosis through monocytes. A.** Experimental design. **B.** Representative pseudocolor plots of MAIT cell staining and quantification from the liver of irradiated female $Ldlr^{-/-CAST}$ and $Ldlr^{-/-CAST}$ $MR1^{-/-}$ mice, which underwent bone marrow transplantation (BMT) receiving bone marrow from female $CCR2^{+/+}$ or $CCR2^{-/-}$ mice, followed by an eight-week high-cholesterol (HC) diet (n = 10 per group). **C.** Representative pseudocolor plots of monocyte staining and quantification from blood of the same irradiated female $Ldlr^{-/-CAST}$ and $Ldlr^{-/-CAST}$ $MR1^{-/-}$ mice after BMT and eight-week of HC diet (n = 10 per group). **D.** Plasma cholesterol levels from irradiated female $Ldlr^{-/-CAST}$ and $Ldlr^{-/-CAST}$ $MR1^{-/-}$ mice receiving bone marrow from female $CCR2^{+/+}$ or $CCR2^{-/-}$ mice followed by an eight-week HC diet (n = 10 per group). **E.** Correlation of cholesterol levels and Oil Red O positive area in the thoracic aorta. **F.** Representative confocal images of the aortic arch stained with DAPI, MOMA-2, Oil Red O and quantification of the MOMA-2 positive area of female $Ldlr^{-/-CAST}$ and $Ldlr^{-/-CAST}$ $MR1^{-/-}$ mice receiving bone marrow from female $CCR2^{+/+}$ (n = 9 per group). Individual data are presented as scatter plots, with the mean and SEM. P-values were determined using the Mann–Whitney test. Dashed lines indicate that the experiments were not done at the same time and the Mann-Whitney test was used for comparison.

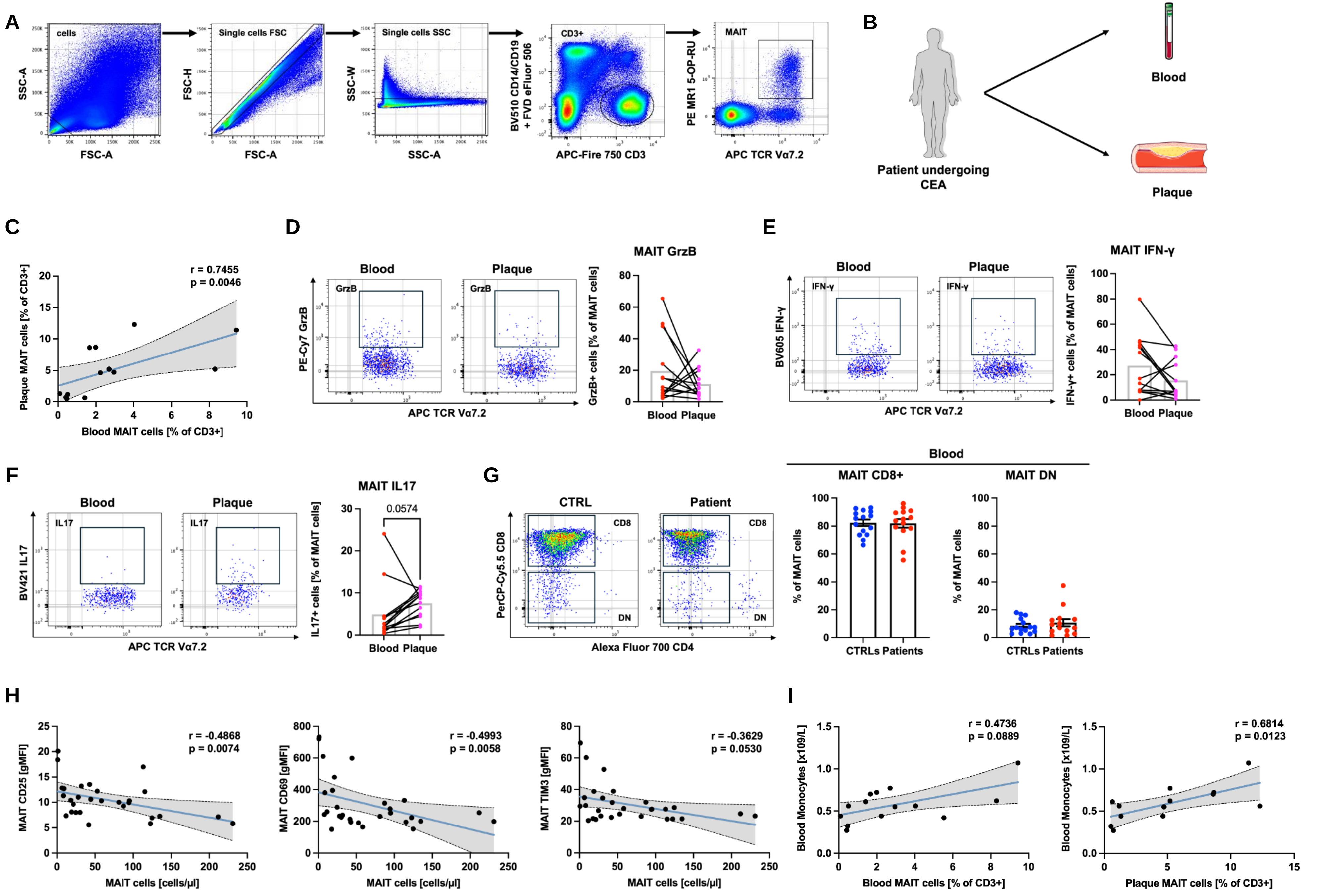

**Supplementary Figure 1**

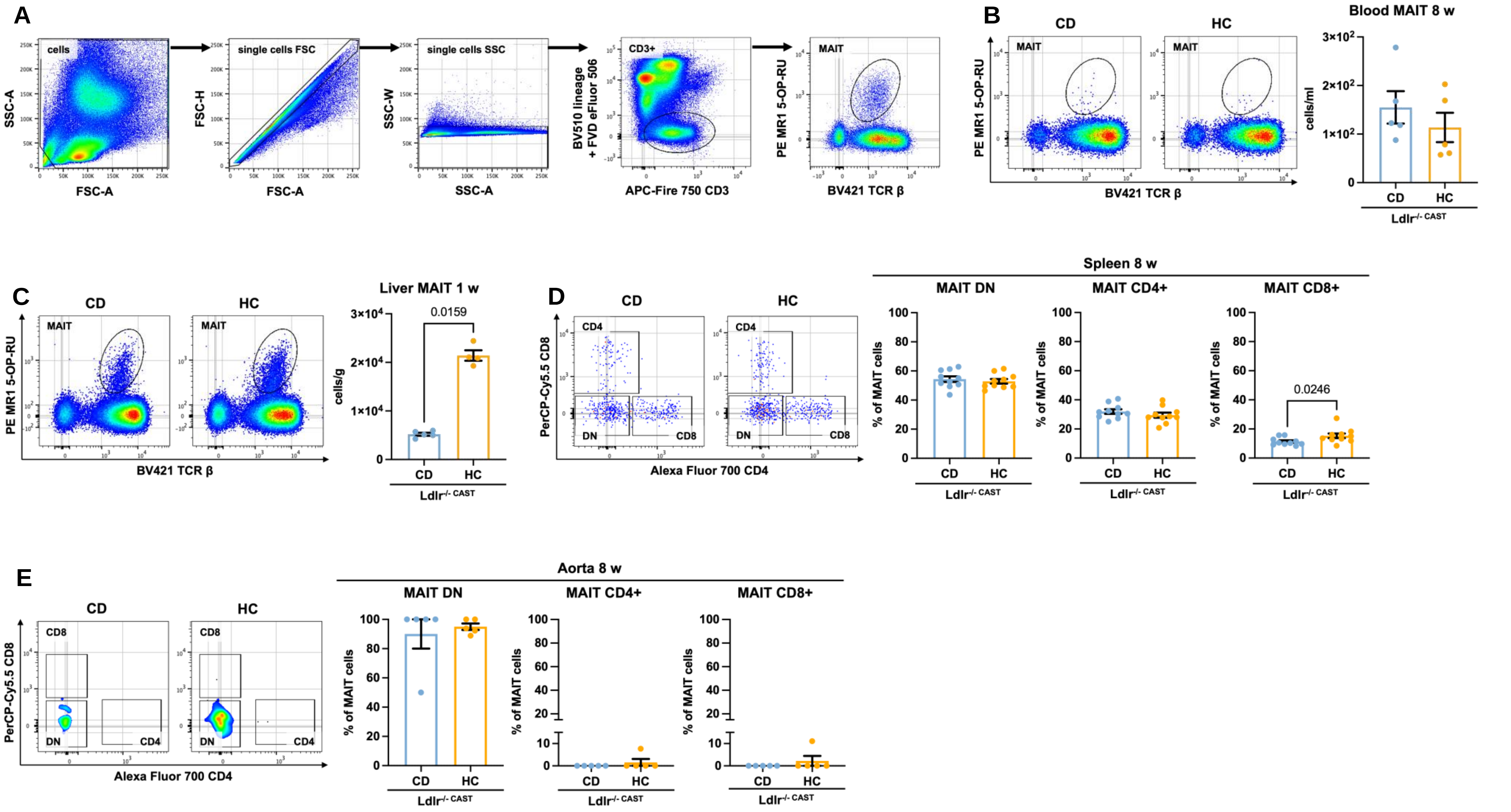

Supplementary Figure 2

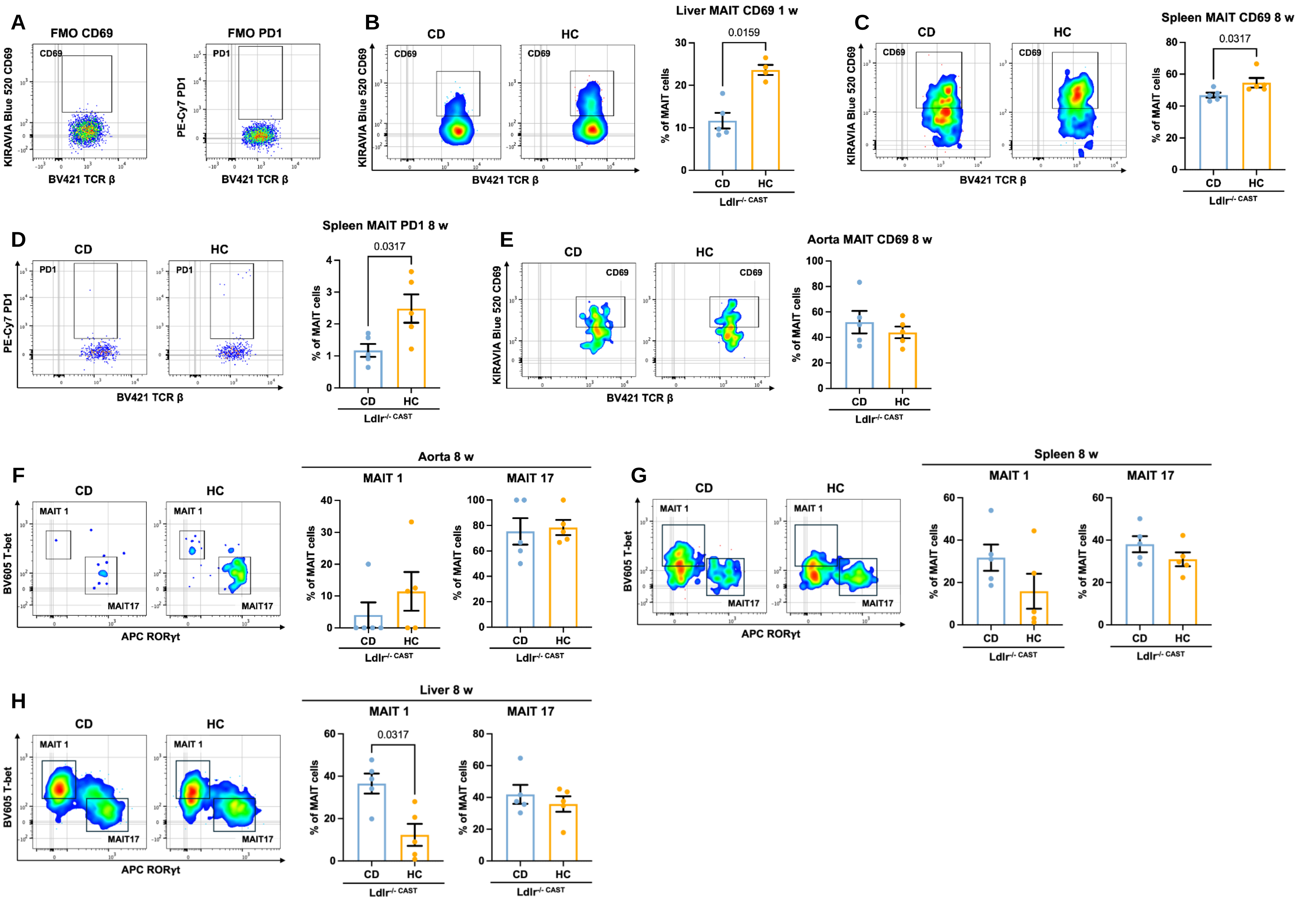

Supplementary Figure 3

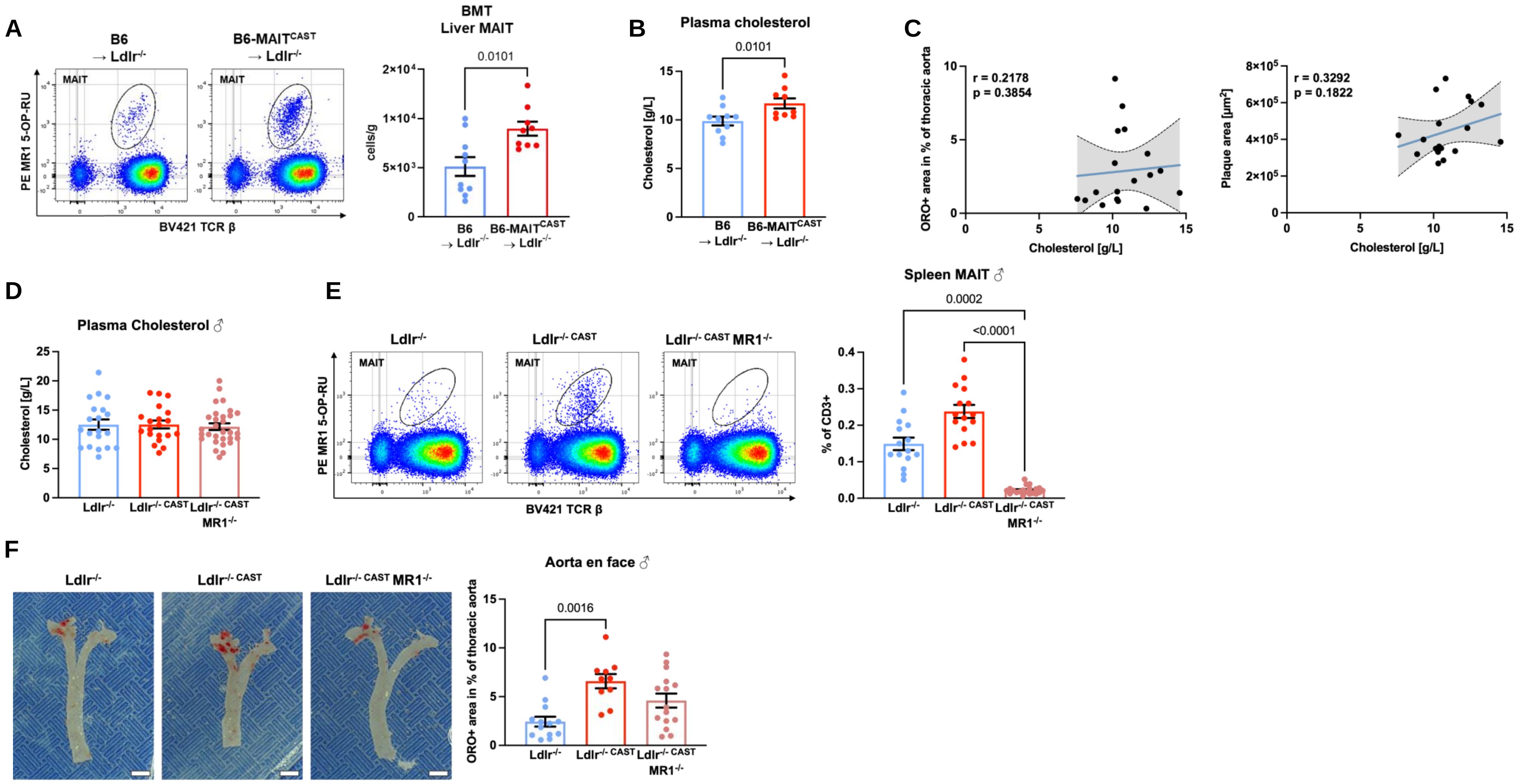

Supplementary Figure 4

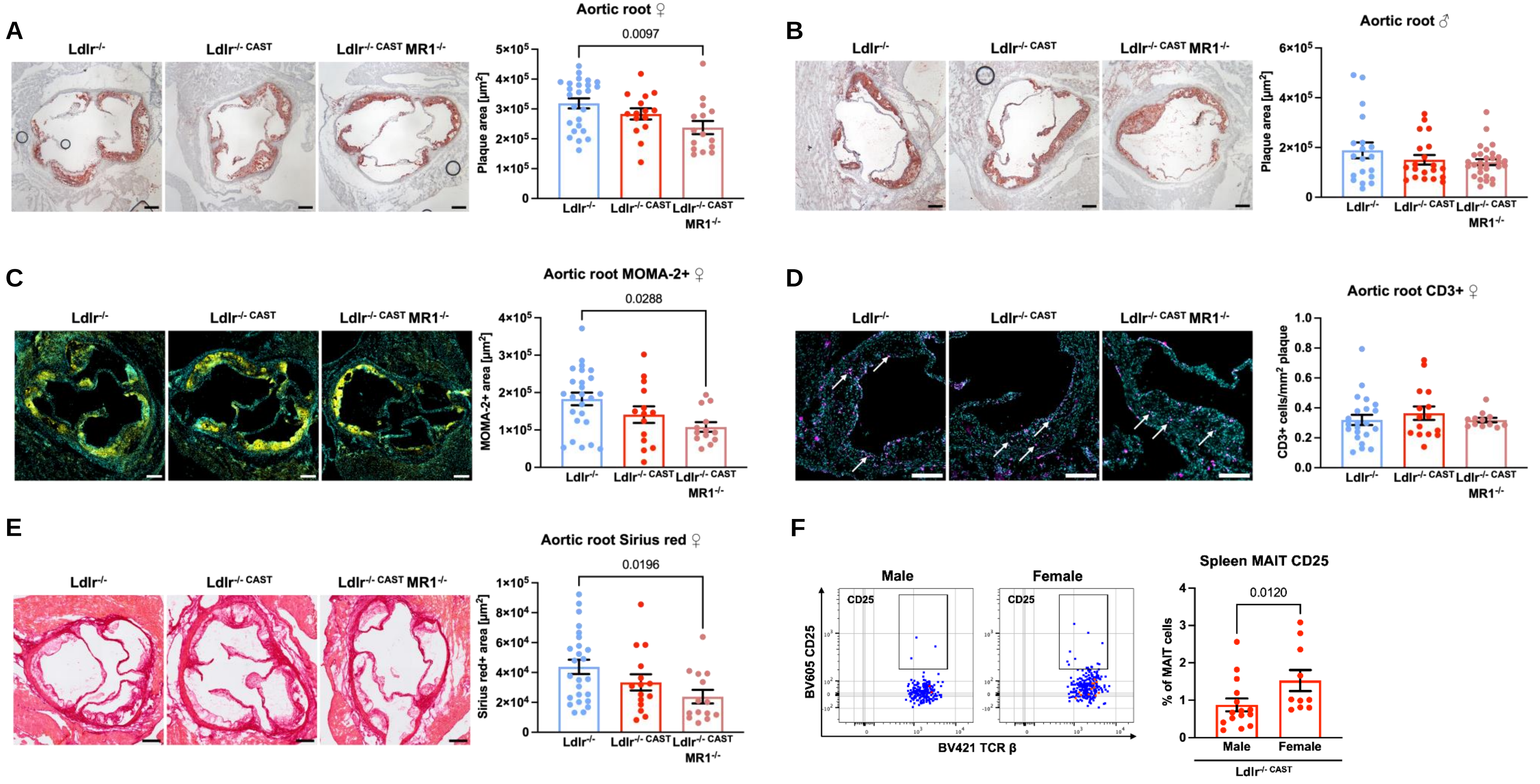

Supplementary Figure 5

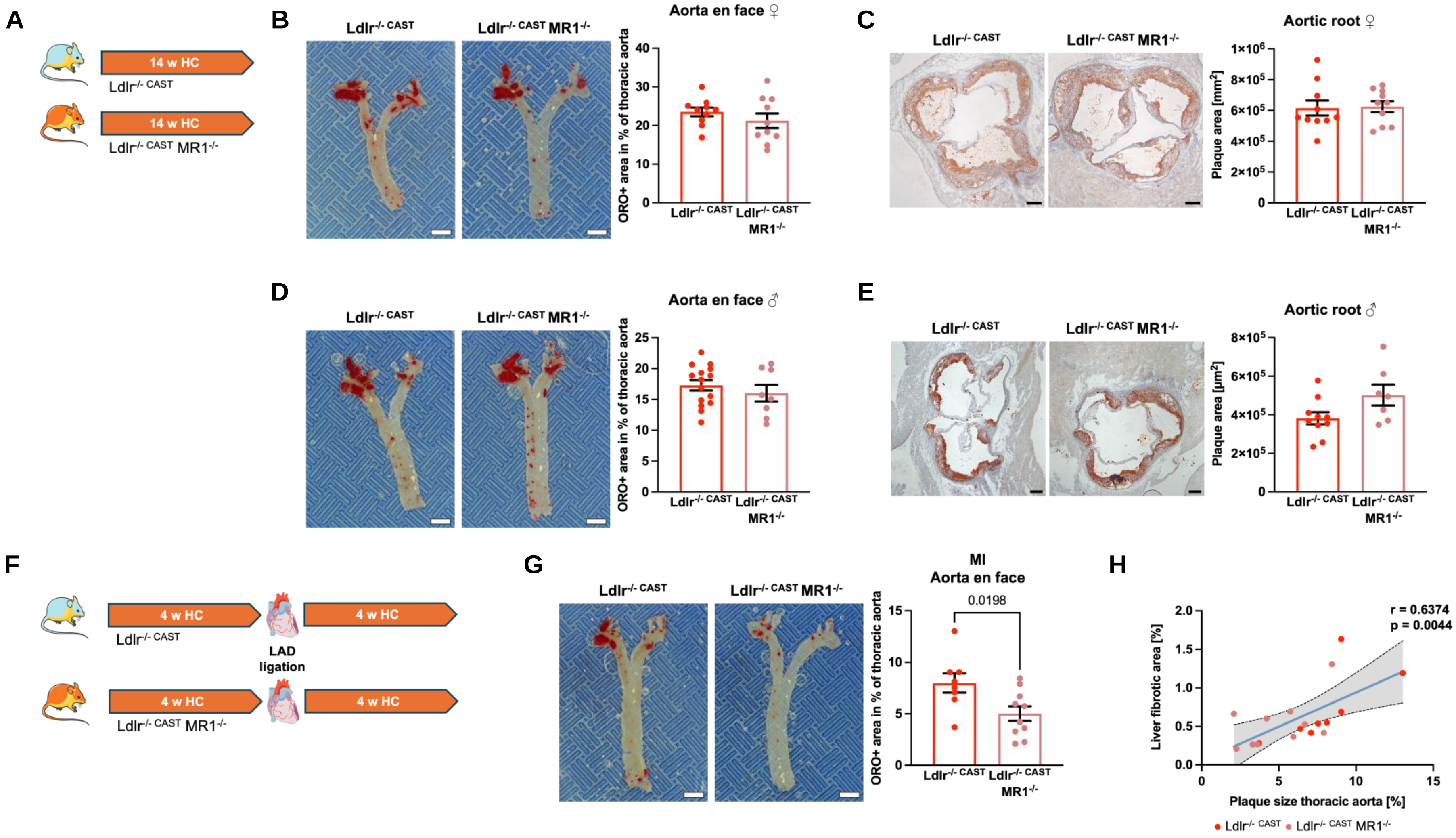

Supplementary Figure 6

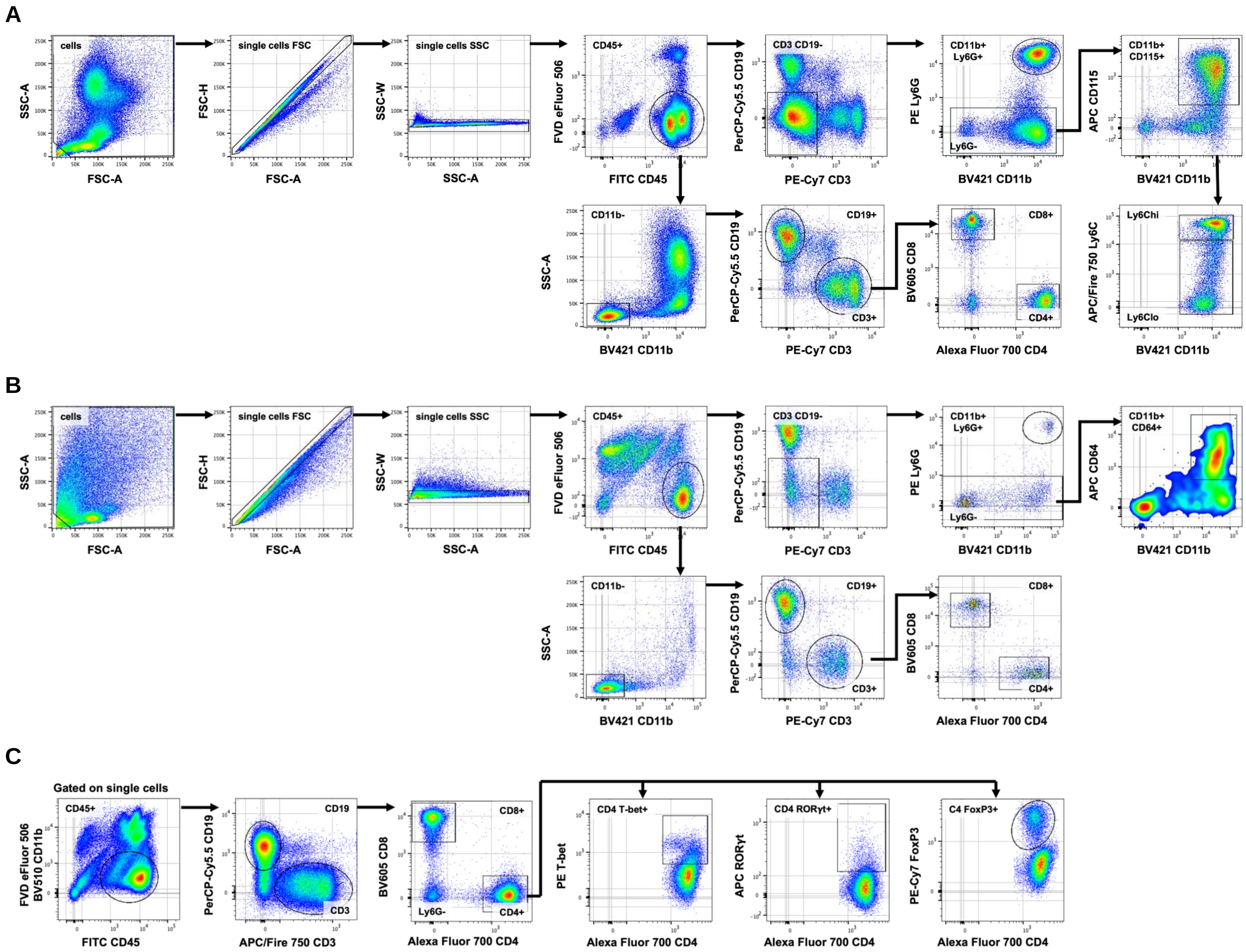

**Supplementary Figure 7**

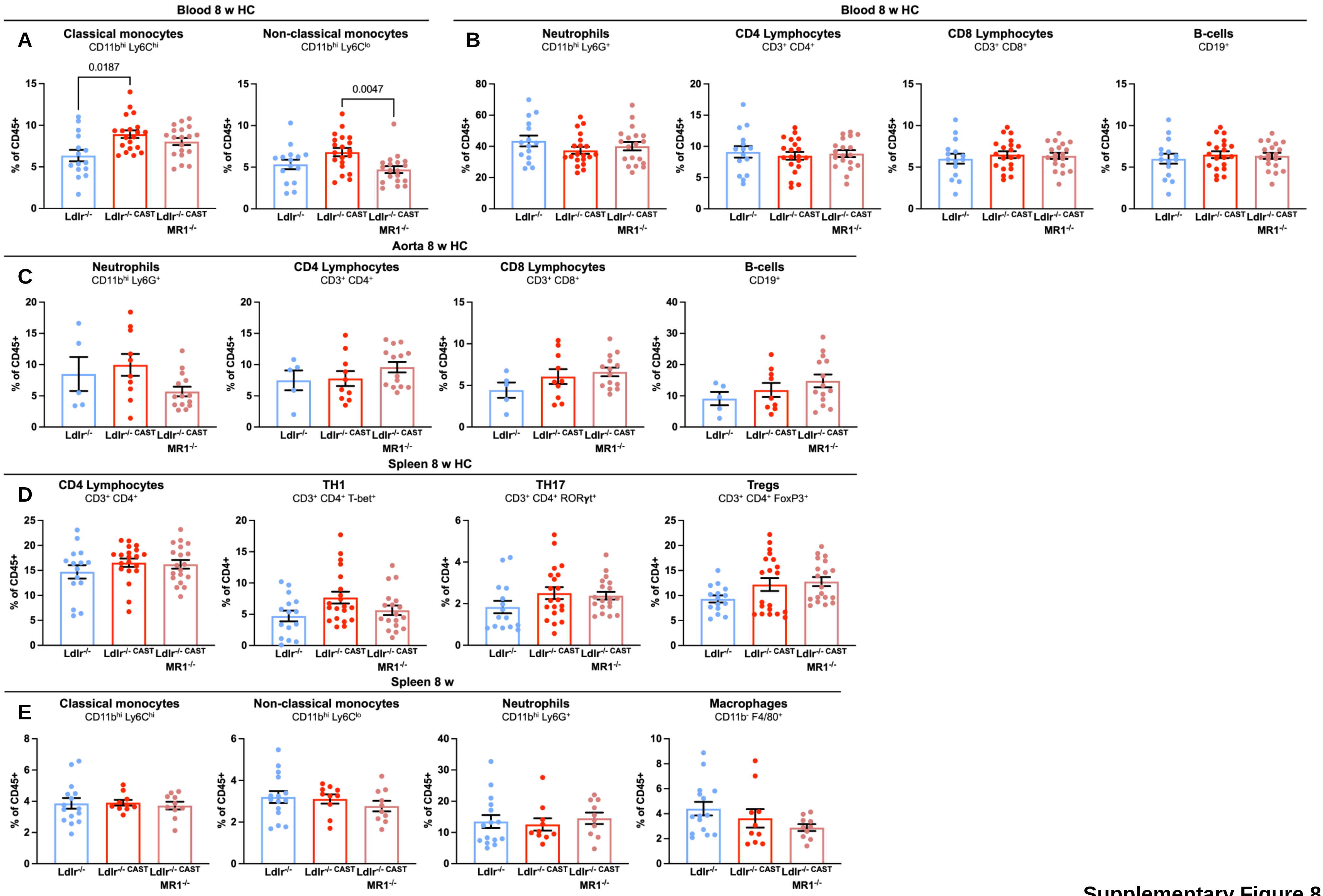

Supplementary Figure 8

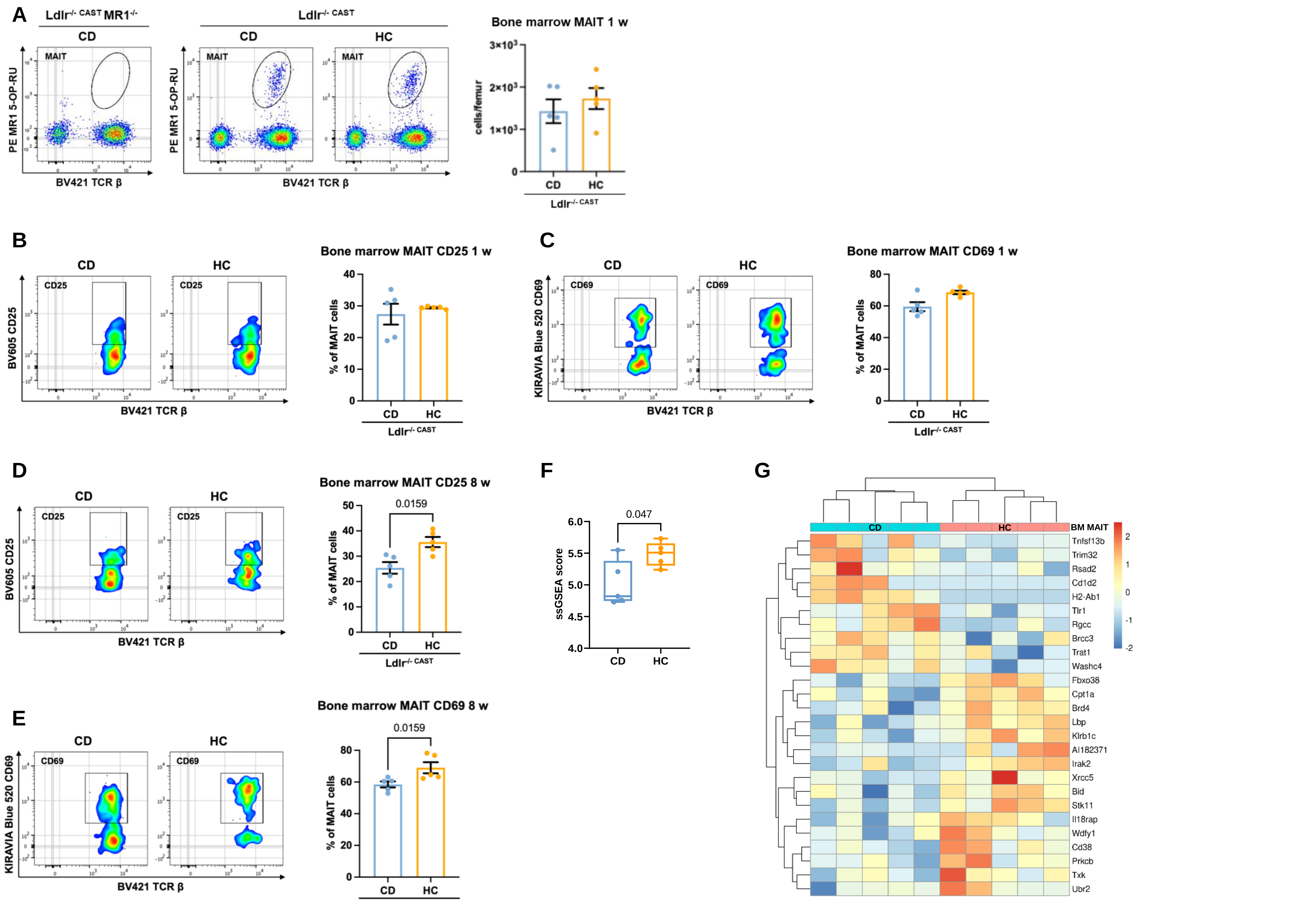

Supplementary Figure 9

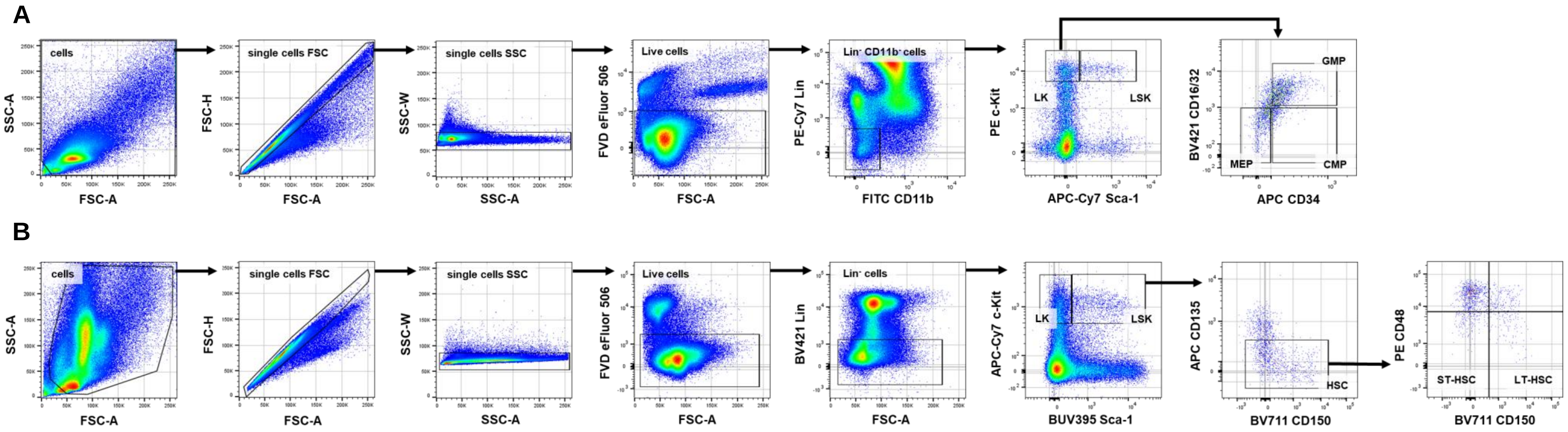

Supplementary Figure 10

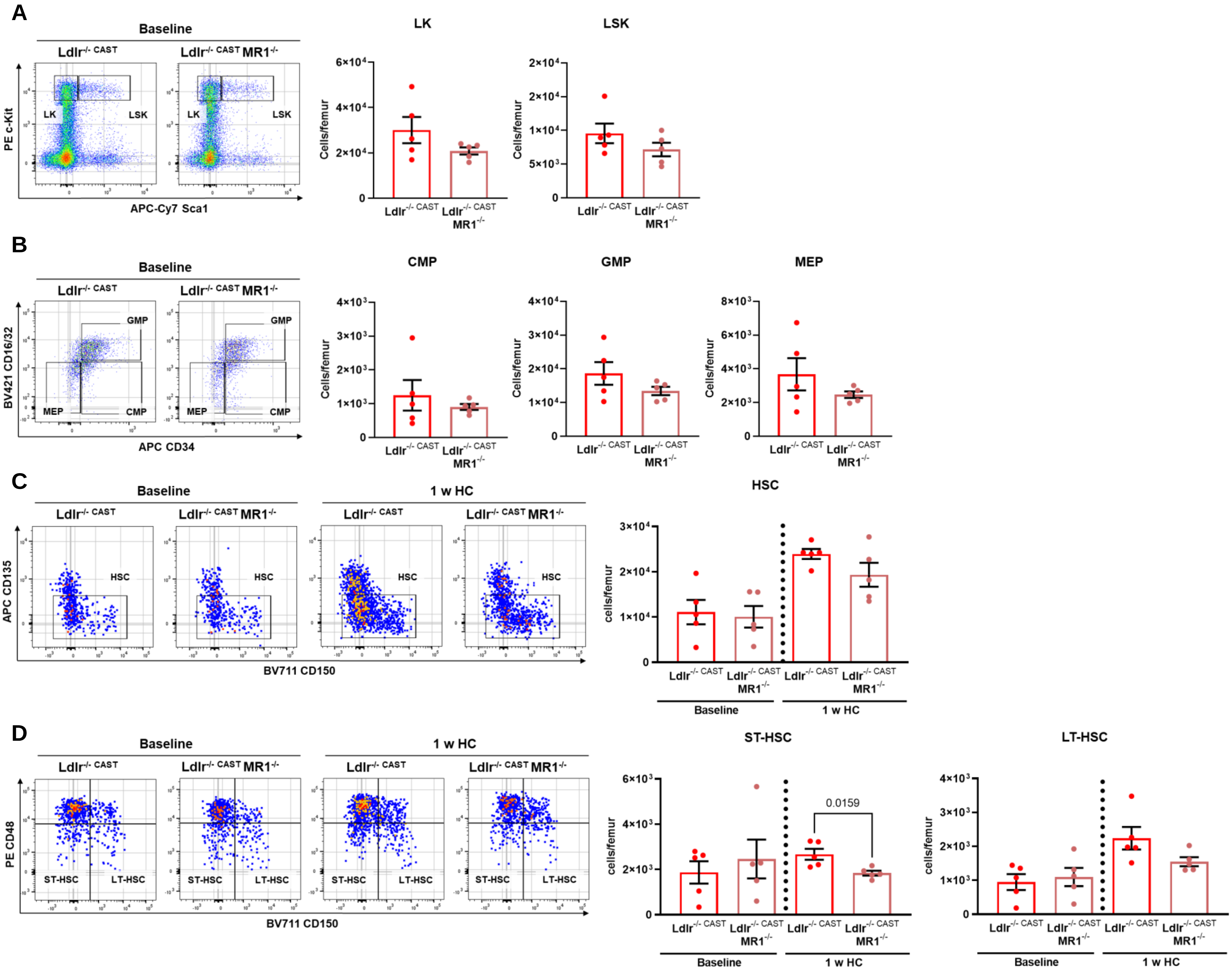

Supplementary Figure 11

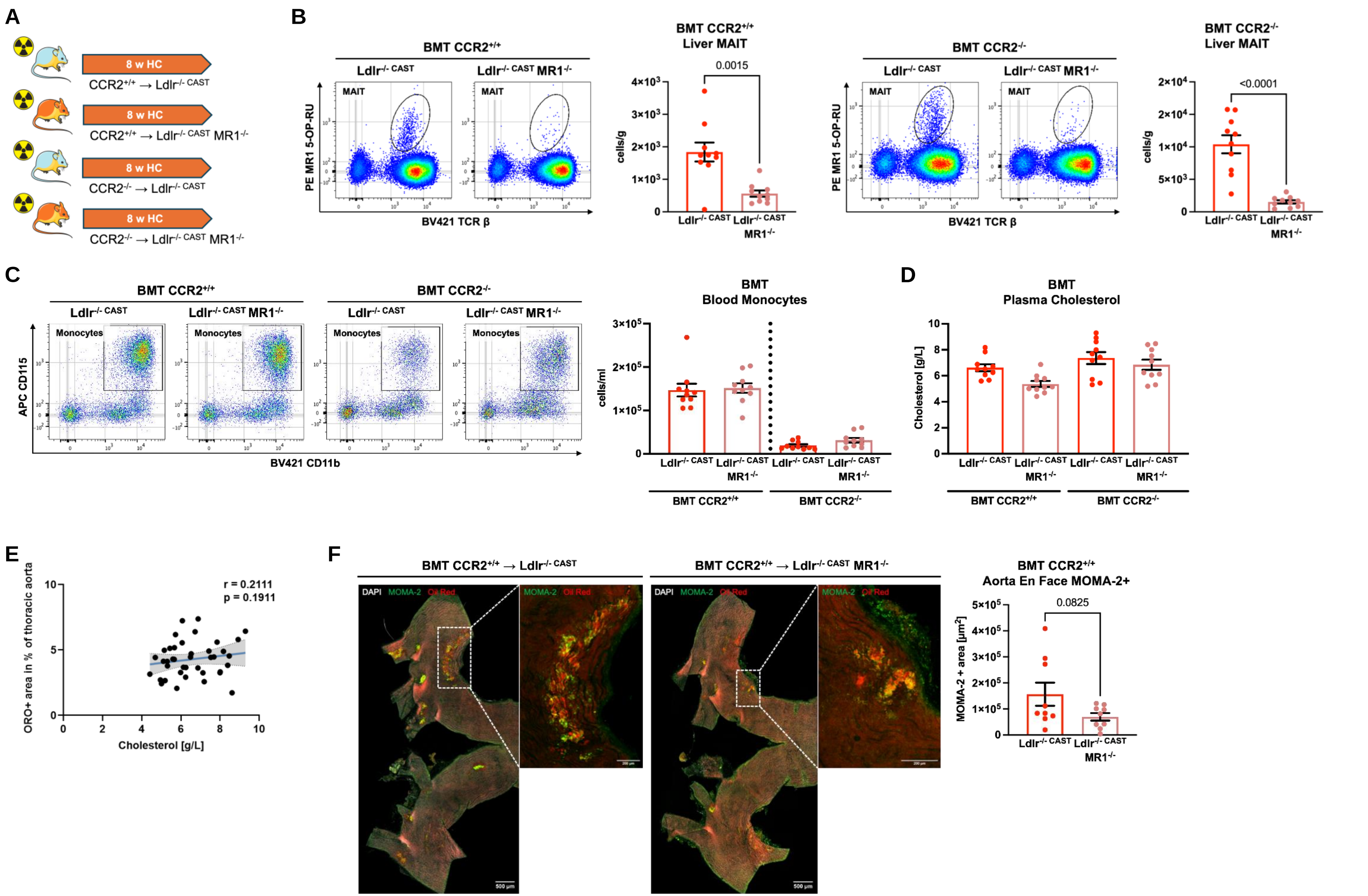

Supplementary Figure 12

Supplementary table 1: Patient's characteristics

| | Patients n = 19 | CTRLs n = 19 |
|---|---|---|
| Male sex (n) | 79 % (15) | 63 % (12) |
| Age (years) | 69 (± 9.5) | 50 (± 19.9) |
| BMI (kg/m2) | 25.2 (± 3.1) | - |
| Smoker (n) | 53 % (10) | - |
| Past smoker (n) | 32 % (6) | - |
| Current smoker (n) | 21 % (4) | - |
| Diabetes mellitus (n) | 11 % (2) | - |
| Hypertension (n) | 74 % (14) | - |
| History of stroke or TIA (n) | 11 % (2) | - |
| History of symptomatic CAD (angina, stenting, CABG, MI) (n) | 16 % (3) | - |
| History of carotid endarterectomy (n) | 5 % (1) | - |
| Peripheral artery disease and AAA (n) | 26 % (5) | - |
| History of cancer (n) | 26 % (5) | - |
| History of atrial fibrillation (n) | 21 % (4) | - |
| eGFR [mL/ min/ 1.73 m2] | 88.0 (± 24.4) | - |
| Total cholesterol [mmol/L] | 4.2 (± 0.8) | - |
| Triglycerides [mmol/L] | 1.4 (± 0.98) | - |
| HDL [mmol/L] | 1.4 (± 0.3) | - |
| LDL [mmol/L] | 2.1 (± 1.0) | - |
| Leukocytes [$x10^9$/l] | 7.00 (± 2.33) | - |
| Monocytes [$x10^9$/l] | 0.61 (± 0.24) | - |
| C-reactive protein [mg/L] | 2.7 (± 5.0) | - |
| Antiplatelets (aspirin or clopidogrel) (n) | 89 % (17) | - |
| Anticoagulants (VKA, NOACs) (n) | 21 % (4) | - |
| Statins (n) | 68 % (13) | - |
| Ezetimibe (n) | 16 % (3) | - |
| ACEi or ARB (n) | 58 % (11) | - |
| Metformin (n) | 11 % (2) | - |
| iSGLT2 (n) | 5 % (1) | - |
| GLP1 analogues (n) | 5 % (1) | - |

Values are median ± SD for continuous variables and percentage (of total count) for categorical data. BMI, body mass index; TIA, transient ischemic attack; CAD, coronary artery disease; CABG, coronary artery bypass graft; MI, myocardial infarction; AAA, abdominal aortic aneurysm; eGFR, estimated glomerular filtration rate; VKA, vitamin K antagonists; NOACs, non-vitamin K antagonist anticoagulant.

**Supplementary Table 1**
